# The last missing piece of the *Triangle of U*: the evolution of the tetraploid *Brassica carinata* genome

**DOI:** 10.1101/2022.01.03.474831

**Authors:** Won Cheol Yim, Mia L. Swain, Dongna Ma, Hong An, Kevin A. Bird, David D. Curdie, Samuel Wang, Hyun Don Ham, Agusto Luzuriaga-Neira, Jay S. Kirkwood, Manhoi Hur, Juan K. Q. Solomon, Jeffrey F. Harper, Dylan K. Kosma, David Alvarez-Ponce, John C. Cushman, Patrick P. Edger, Annaliese S. Mason, J. Chris Pires, Haibao Tang, Xingtan Zhang

## Abstract

Ethiopian mustard (*Brassica carinata*) is an ancient crop with significant potential for expanded cultivation as a biodiesel feedstock. The remarkable stress resilience of *B. carinata* and desirable seed fatty acid profile addresses the ongoing food vs. fuel debate as the crop is productive on marginal lands otherwise not suitable for even closely related species. *B. carinata* is one of six key *Brassica spp*. that share three major genomes: three diploid species (AA, BB, CC) that spontaneously hybridized in a pairwise manner, forming three allotetraploid species (AABB, AACC, and BBCC). Each of these genomes has been researched extensively, except for that of *B. carinata*. In the present study, we report a high-quality, 1.31 Gbp genome with 156.9-fold sequencing coverage for *B. carinata* var. Gomenzer, completing and confirming the classic *Triangle of U*, a theory of the evolutionary relationships among these six species that arose almost a century ago. Our assembly provides insights into the genomic features that give rise to *B. carinata*’s superior agronomic traits for developing more climate-resilient *Brassica* crops with excellent oil production. Notably, we identified an expansion of transcription factor networks and agronomically-important gene families. Completing the *Triangle of U* comparative genomics platform allowed us to examine the dynamics of polyploid evolution and the role of subgenome dominance in domestication and agronomical improvement.

## Introduction

*B. carinata* has been traditionally cultivated as a dual oilseed and leafy vegetable crop in the Ethiopian highlands as a staple food in rural areas, especially during times of food shortages (Ojiewo et al., 2013). Locally referred to as gomenzer, *B. carinata* is among the oldest Ethiopian crops and has recently gained traction as an alternative biofuel feedstock in other Mediterranean climates around the world, such as Canada, India, and Italy, for its adaptability and notable productivity on marginal land with low input (Taylor et al., 2010; Cardone et al., 2003). According to the classic *Triangle of U* model, *B. carinata* spontaneously arose from the hybridization between two progenitor species, *B. nigra* and *B. oleracea* (U, 1935). The *Triangle of U* describes the evolutionary relationships among six globally important *Brassica* species that share independently-evolved versions of three core genomes, termed A, B, and C (Figure 1). Three diploid species, *B. rapa* (ArAr, *2n =* 20), *B. nigra* (BnBn, *2n =* 16), and *B. oleracea* (CoCo, *2n* = 18), participated in pairwise hybridizations, forming three allotetraploids: *B. juncea* (AjAjBjBj, *2n =* 36), *B. napus* (AnAnCnCn, *2n =* 38), and *B. carinata* (BcBcCcCc, *2n =* 34). The overlapping nature of the genomic relationships among these six *Brassica spp.* allows the evolution of a given (sub)genome to be compared in different genomic environments.

**Figure 1.**
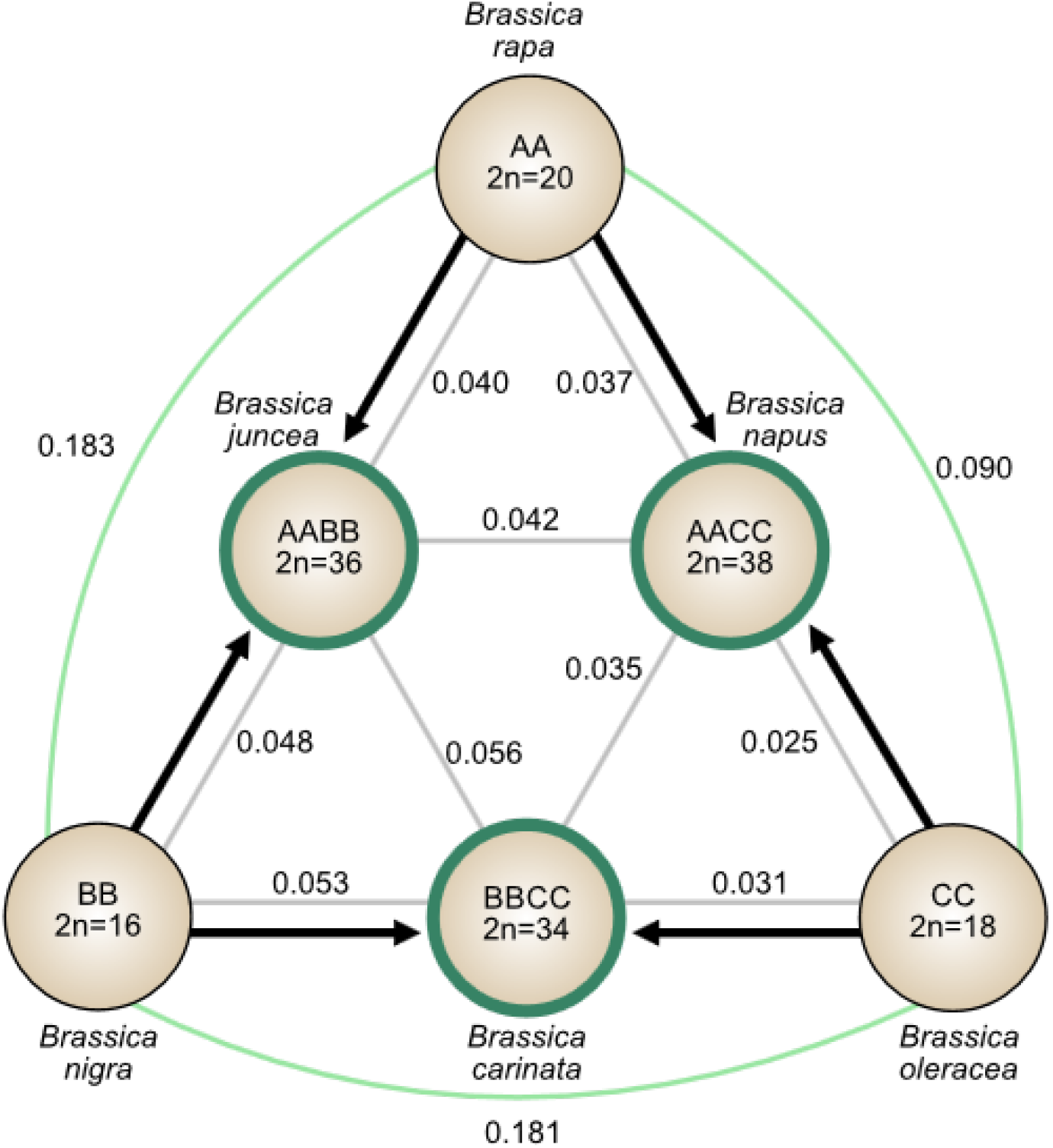
Genetic relationships between *Brassica* species in the *Triangle of U* model. The *Triangle of U* describes the genetic relationships among the *Brassica* species that share the same three core genomes. The three diploid species (*Brassica rapa*, *B. nigra*, and *B. oleracea*) representing the AA, BB, and CC genomes, respectively, are shown at the nodes of the triangle. The arrows along the sides of the triangle indicate the pairwise hybridization events that occurred to generate the three allotetraploid species (*B. juncea*, *B. napus*, and *B. carinata*) highlighted in green. The synonymous substitution per site (*K*_s_) values between each species are shown to convey the degree of differentiation of the shared genomes (subgenomes). Larger *K*_s_ values indicate that the subgenomes have diverged more over evolutionary time.

**Figure 2.**
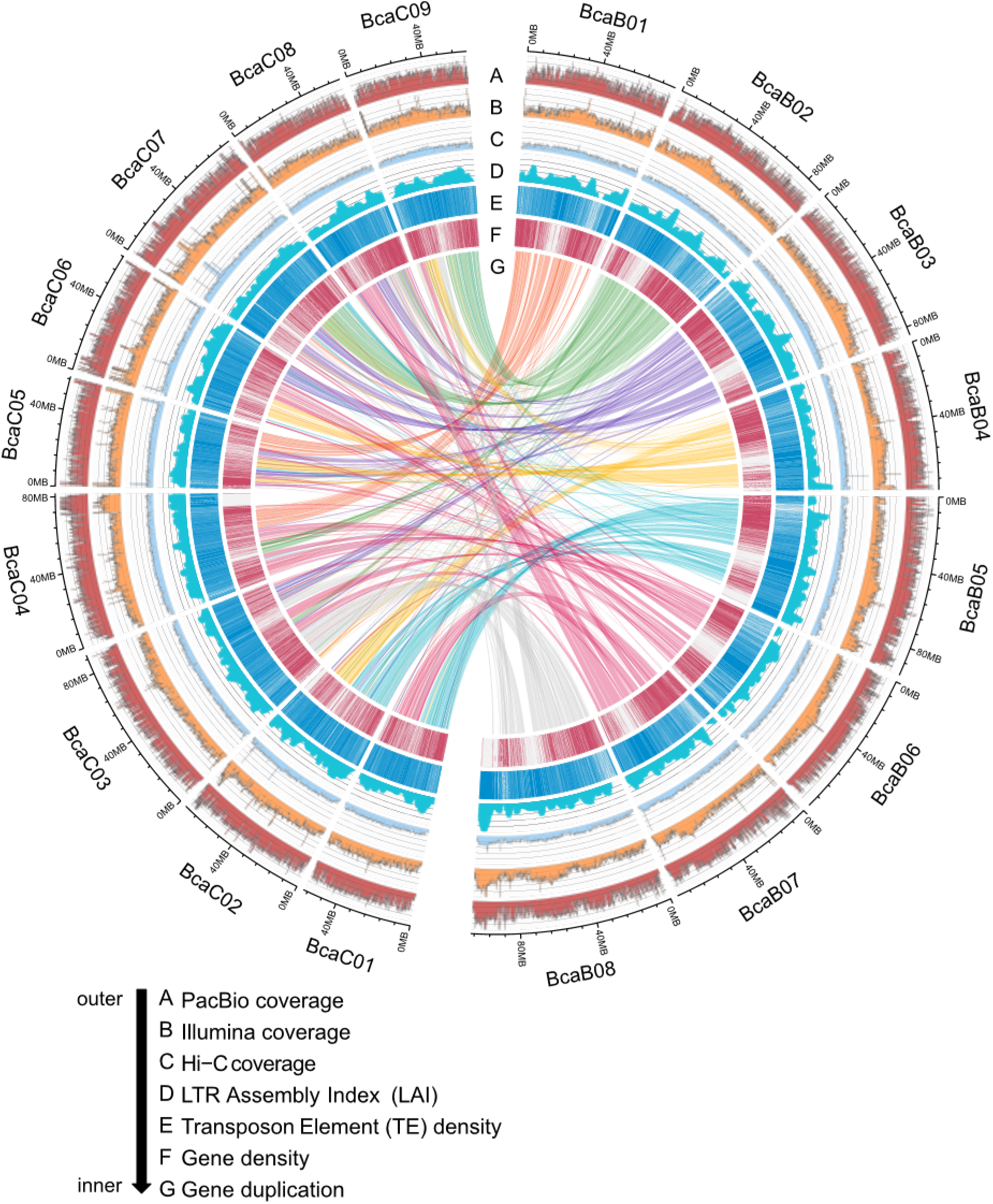
Chromosomal features of the *Brassica carinata* Gomenzer genome assembly. The *B. carinata* genome comprises 17 total chromosomes with eight chromosomes belonging to the Bc subgenome (BcaB01-BcaB08, right-hand semicircle) and nine chromosomes belonging to the Cc subgenome (BcaC01-BcaC09, left-hand semicircle). The size of each chromosome is scaled according to its assembled length. Homoeologous relationships between the two subgenomes are indicated with links connecting Bc subgenome with Cc subgenome homoeologs. The tracks, from outermost to innermost, display the sequences depths for PacBio, Illumina, and Hi-C data, long terminal repeat (LTR) assembly index (LAI), transposable element (TE) density, gene density, and gene duplication. All the plots are drawn in a 100-Kb sliding window.

This ability to investigate the evolutionary trajectories of the same core genome in different species is ideal for elucidating the mechanisms that act on duplicated genes over the diploidization process, such as functional divergence of gene families and the biased fractionation of one parental genome, or subgenome. Furthermore, the precedence of these *Brassica spp.* as a global vegetable source, including crops such as canola, broccoli, cabbage, and mustard seeds, adds economic value to the elucidation of the genomic features that contribute to their trait variations. Although the *Triangle of U* was introduced in 1935, the utility of this well-established comparative genomics platform was not truly harnessed until the large, highly syntenic, allotetraploid genomes could be assembled at the chromosome scale. While all angiosperms evolved from ancient polyploids, polyploidy is particularly common among major crops with domesticated plants having more polyploidization events in their evolutionary history than wild species (Salman-Minkov et al., 2016; Wendel, 2015). As polyploidization and crop domestication are closely associated, shedding more light on the genetic consequences of polyploidization is essential for understanding crop domestication, and on a larger scale, angiosperm evolution.

Here we generated a complete and comprehensive characterization of the *B. carinata* genome to help illustrate the genetic basis of its numerous profitable traits. *B. carinata* is a highly productive mustard and one of the most drought and heat tolerant species within the economically important Brassicaceae family (Kumar et al., 1984; Malik, 1990). Additionally, the species’ cold tolerance allows its use in winter fallow fields, reducing nutrient and sediment run-off to preserve the surrounding environment’s water quality (Hoghooghi et al., 2020). The seed oil has a large erucic acid proportion (C22:1), which is toxic to humans but ideal for biodiesel production due to the resulting high cetane number and oxidative stability (Cardone et al., 2003; Folayan et al., 2019). Thus, the high productivity under low input conditions and high erucic acid content make *B. carinata* a more sustainable alternative to the current biodiesel feedstocks.

Approximately 50% of the world’s land area is already under arid, semi-arid, or dry sub-humid conditions that restrict the yield of major crops, an issue that is exacerbated by global climate change (Zika and Erb, 2009; Lobell et al., 2011; Read et al., 2013). Biofuels could offer sustainable energy security while decreasing greenhouse gas emissions; however, more efficient crop and land use are also necessary to reap such climate benefits (Kazamia and Smith, 2014; Searchinger et al., 2018). Thus, there is a clear and present need to adopt more climate-resilient crops into our bioenergy feedstock portfolio that can use limited freshwater resources more efficiently and make better use of marginal or abandoned lands in semi-arid and tropical sub-humid regions of the Earth (Husen et al., 2014). Furthermore, oilseed *Brassica* lines carrying advantageous alleles at the genomic regions defined here could be useful as introgression donors in breeding for improved climate resilience and oil production, which could decrease the costs of renewable diesel and jet fuels (Wei et al., 2016; Zhang et al., 2017; Basili and Rossi, 2018).

## Results

### Construction of our high-quality *B. carinata* genome provides insight into asymmetric distributions in genomic features of the Bc and Cc subgenomes

We assembled the *B. carinata* A. Braun var. Gomenzer genome (USDA, PI 273640) using PacBio, Illumina, and Hi-C sequencing data. The initial PacBio read-based assembly produced a 1.31 Gbp draft genome with a contig N50 value of 602.9 kb and 156.9-fold coverage from 140.76 Gbp of PacBio long reads and 63.3 Gbp of Illumina short reads (Table S2-S3, Figures S2-S3). The 11,531 contigs assembled from the PacBio sequences were segregated according to their subgenome, Bc or Cc, using the *B. nigra* (Bn) and *B. oleracea* (Co) genomes as the reference (Liu et al., 2014; Wang et al., 2019). The subgenome-clustered contigs were then ordered and oriented into scaffolds according to the Hi-C loci contact matrix, anchoring the scaffolded sequence into the expected 17 pseudo-chromosomes (Figure S5C, Table S4,). The scaffold N50 of the final assembly increased to 78.8 Mbp (Table S6). The final 1.31 Gbp assembly covered a 95% confidence interval of the total estimated genome size of 1.28 Gbp by flow cytometry and 1.31 Gbp by *k*-mer analysis (Figure S5A). The complete genome assembly was more extensive than those of the other two allotetraploid species *B. juncea* (AjBj, ∼955 Mbp) and *B. napus* (AnCn, ∼976 Mbp), making the *B. carinata* genome the largest among the *Triangle of U* (Table S7) (Liu et al., 2014; Chalhoub et al., 2014; Yang et al., 2016; Perumal et al., 2020). Comparison of the assembled subgenomes revealed an asymmetry in size. The Bc subgenome (686.6 Mbp) is 62.7 Mbp larger than the Cc subgenome (623.9 Mbp, Table S6). As with total genome size, both subgenomes are larger than the assemblies of their respective progenitor genomes (Bn at ∼608 Mbp and Co at ∼555 Mbp).

In line with the larger genome size, the *B. carinata* genome has the largest gene content of the *Triangle of U*, containing 119,303 annotated gene models predicted to generate 197,479 transcripts (Figure 3A, Table S13). We also identified asymmetry in the total gene content between the two subgenomes, with the larger Bc subgenome containing 54.3% of the total gene content (64,787 vs. 54,516, Table S13). The larger Bc subgenome is more gene rich, with exons comprising 8.6% of the subgenome compared to the 6.8% exon content its counterpart. While genomic rearrangements inherent to the hybridization process are likely factors, the observed size asymmetry could also be affected by the genome size of the donor genotype. Recent evidence suggests that the variations in genome size between accessions have biological relevance and are not just an effect of variations in genome quality. For example, a comparative study of the *B. rapa* Z1 and Chiifu genotypes cytogenetically validated several large chromosomal variants, which together contributed to a 16% difference in genome size between them (Boutte et al., 2020). Analysis of additional genotypes, construction of a *B. carinata* pan-genome, and production of linkage mapping populations will aid in identifying the *B. nigra* (Bn) and *B.oleracea* (Co) donor genotypes.

**Figure 3.**
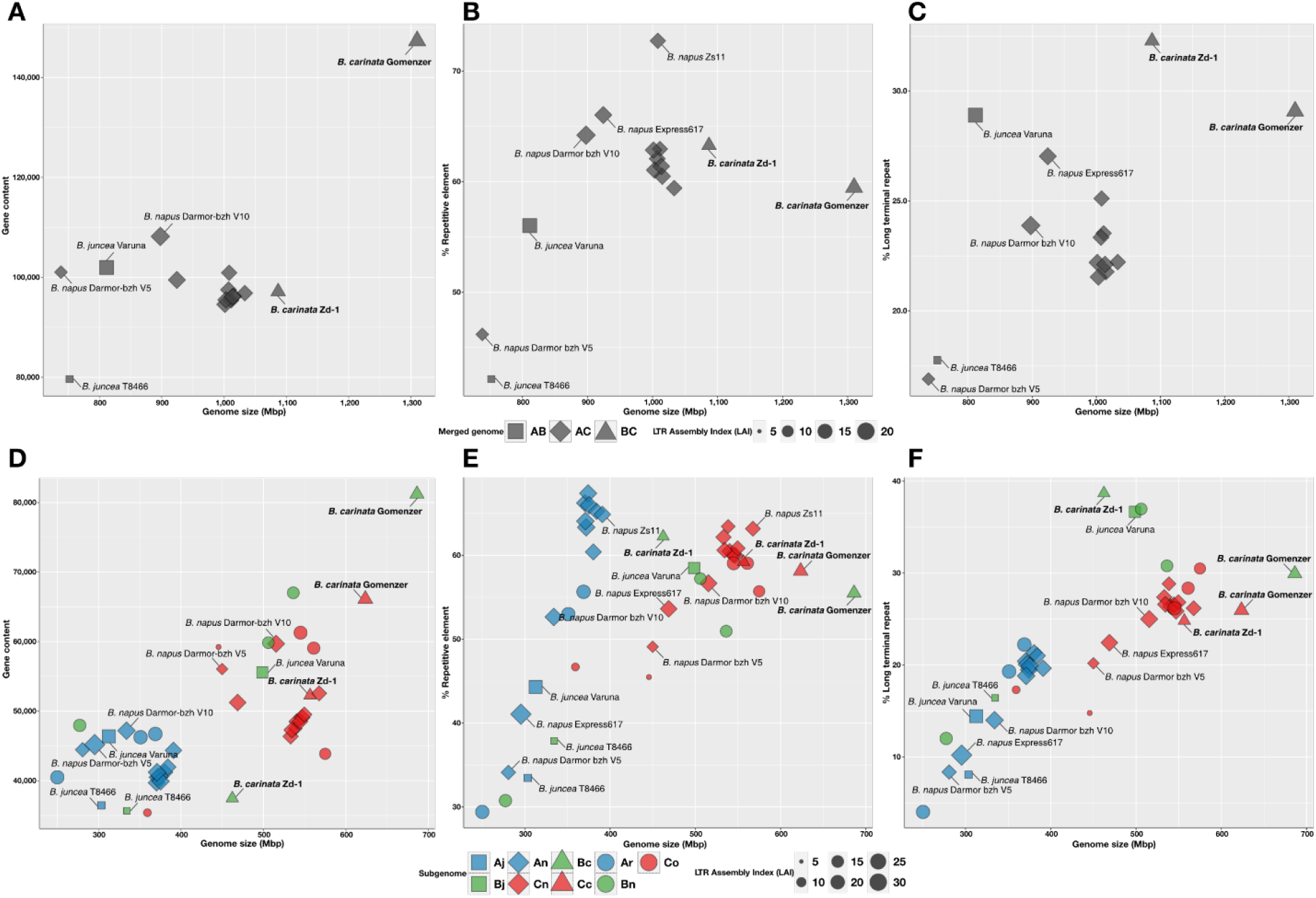
Dot plot comparisons of genome size and genetic features among assembled *Brassica* genomes. The upper panels (A, B, and C) compare whole genome assemblies of the *Triangle of U* tetraploids and the lower panels (D, E, and F) compare subgenome assemblies among all six species. The x-axes represent the sizes of the *Brassica* genome assemblies in Mbp, and the y-axes show either gene content (left), repetitive element content (middle), or long terminal repeat (LTR) content (right). The shape of the data points indicates the genome and the sizes of the data points are scaled according to the assembly’s LTR assembly index (LAI), with larger LAI values indicating a higher quality assembly. For the bottom panels, the three subgenomes are assigned as blue (A), green (B), or red (C). The three allotetraploid species are represented as unique shapes: square (*B. juncea*), diamond (*B. napus*), and triangle (*B. carinata*), with their progenitor species (*B. rapa*, *B. nigra*, and *B oleracea*) represented as circles. The *B. carinata* assemblies, Gomenzer and Zd-1, are highlighted in bold, and other assemblies of interest are marked on each panel.

**Figure 4.**
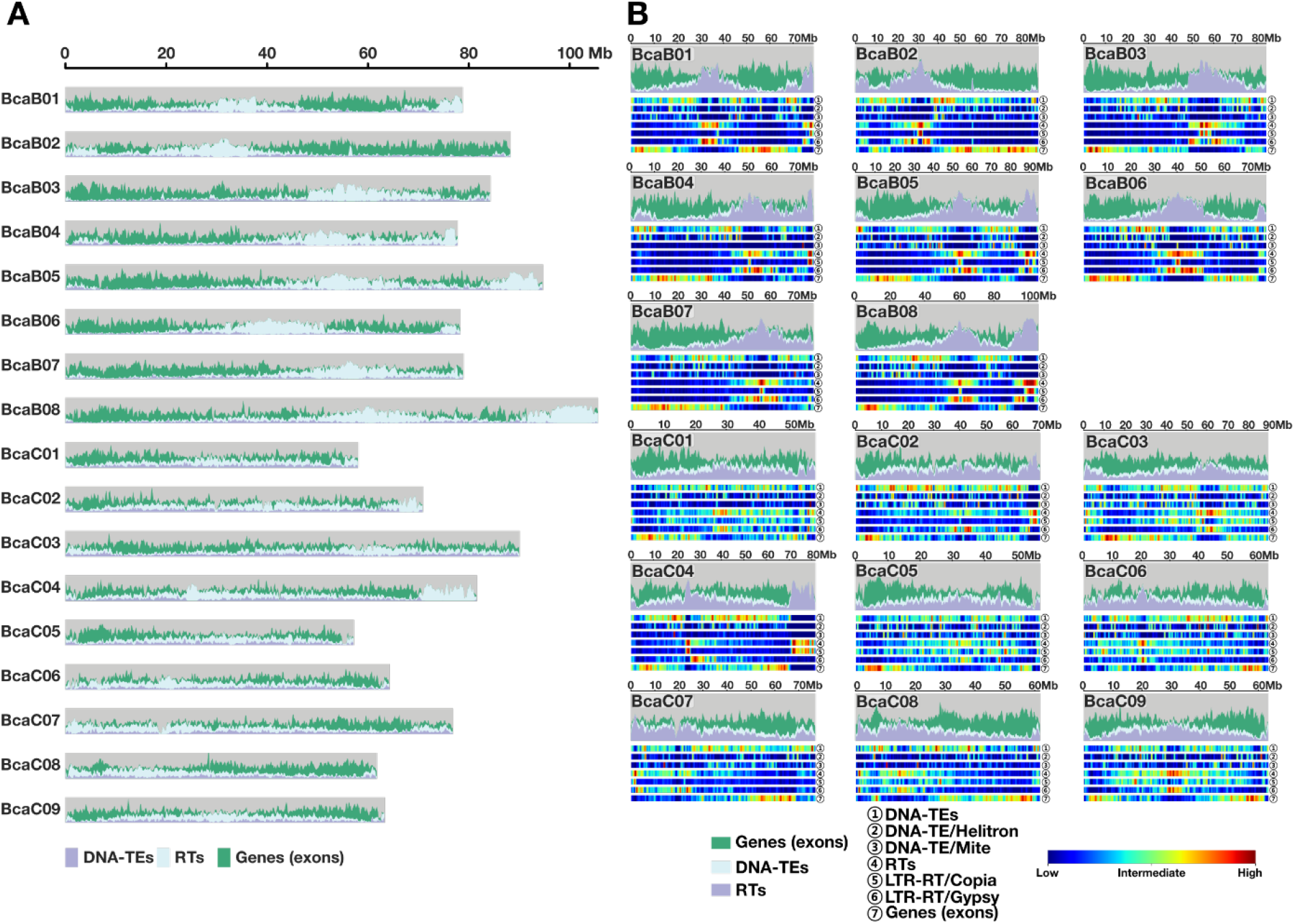
Genomic landscape of *Brassica carinata* chromosomes. **(A)** Distribution of genomic features. The area chart indicates the proportion of class II DNA transposons (DNA-TEs) in purple, Retrotransposons (RTs) in light blue, and exons in green. The x-axis represents chromosome length and the y-axis represents each of the 17 chromosomes. **(B)** Detailed genomic materials are described along with *Brassica carinata* 17 subgenomes. Gene exons in green, DNA-TEs in light blue, and RTs in purple are distributed in the area chart. The heat map tracks represent DNA-TEs, LTR-RTs, and exons for each chromosome, ranging from blue to red with increasing frequency per window size.

The direct relationship between genome size and gene content was also identified in the other two *Triangle of U* allotetraploids. A direct relationship was found in *B. napus* (AnCn), with the dominant Cc subgenome containing ∼54% of the annotated gene and contributing to ∼59% of the genome size (Figure 3D) (Bird et al., 2020). Similarly, the *B. juncea* (AjBj) dominant Bj subgenome is also larger, comprising ∼57% of the total genome and ∼52% of the gene content (Table S13). To investigate whether these asymmetries are inherent to the *Triangle of U* core genomes, we found that the A (sub)genomes generally have a smaller genome size and lower gene content. Conversely, the C (sub)genomes generally have a larger genome size along with a higher gene content. However, the B (sub)genomes are much more variable in terms of genome size and gene content among the six species (Figure 3D-F).

A survey of BUSCOs (Benchmarking Universal Single-Copy Orthologs) was used to assess the completeness of our genome assembly (Simão et al., 2015). A total of 412 of the 425 (96.9%) conserved genes in the Viridiplantae single-copy ortholog dataset were identified in the *B. carinata* genome, 84% of which were duplicated (Figure S9). Identifying the majority of BUSCOs in a genome indicates a high-quality assembly, and the magnitude of duplications reflects the numerous duplications the *B. carinata* genome has experienced. Our RNA-Seq analysis resulted in low rates (∼48%) of assignment of expressed sequences to the chromosomes, likely due to the ambiguity in read mapping caused by the high homology between the two subgenomes (Figure S7C). These results denote apparent gene redundancy in *B. carinata* similar to that observed for many other paleopolyploidization events in plants. Our results also strengthen the hypothesis that gene retention is related to plant species survival in changing environments (Fawcett et al., 2009; Lohaus and Van de Peer, 2016) and to the maintenance of proper relative gene dosage for the function of macromolecular complexes and signaling pathways (Birchler and Veitia, 2012).

### Transposable element propagation patterns facilitate the increased size of the *B. carinata* genome and gene family expansions

Our repeat analysis identified 192 transposable element (TE) classes comprising 62.5% (818.6 Mbp) of the whole genome (Table S9). As with genome size, we found a significantly higher TE content for both subgenomes compared to the respective B and C genomes of the related *Brassica* species (Bj, Bn, Cn, and Co, Table S10). The prevalent TE content indicates that the larger genome size of *B. carinata* might have resulted from the evolution and propagation of TEs (Peterson et al., 2002; Kidwell, 2002; Canapa et al., 2015). Mechanisms that eliminate repetitive sequences, such as illegitimate recombination, have been previously implicated as an avenue of subgenome evolution in *B. rapa* (Ar) (Tang et al., 2012), yet appear to be less active in *B. carinata*.

An extensive genome-wide subgenome comparison identified an asymmetry in TE content between the two subgenomes. The smaller Cc subgenome is more TE dense, with TEs comprising 65.2% of the subgenome, compared to the 60.0% TE content of the Bc subgenome (Table S10). The asymmetric TE distribution might be due to differences in the progenitor genomes, with the *B. nigra* (Bn) and *B. oleracea* (Co) genomes having a 51.9% and 64.0% TE content, respectively. The Bc subgenome also has less TEs per gene (16.7) compared to the Cc subgenome (18.0). The ratio of genes to TEs is reflected in the progenitor species’ genomes with *B. nigra* (Bn) having a considerably lighter TE load (9.88) than *B. oleracea* (Co, 17.4, Table S10, S13). A lower TE load in one progenitor genome is hypothesized to determine dominance of that subgenome due to differences in methylation status around genes (Cheng et al., 2018). With no significant differences in TE dynamics noted, the asymmetry in TE content is likely due to Bc biased TE fractionation.

In general, genome size increases with raising trend of repetitive elements (Blommaert et al., 2019). Assuming that increased genome size is related to repeat content, we would expect a lower exon content based on previous reports (Schubert and Vu, 2016). The results of our TE analysis, coupled with those of our exon analysis, confirms a negative correlation between exon content and repeat content in the *B. carinata* subgenomes, which was supported by our data for gene and repeat distributions in euchromatin and heterochromatin (Figure 5) (Bowers et al., 2005). However, the *B. juncea* (AjBj) subgenome sizes that positively correlate with repetitive elements and gene content. The larger, more gene-rich Bj subgenome contains a higher proportion of repetitive sequence than the Aj subgenome (Figure 3).

**Figure 5.**
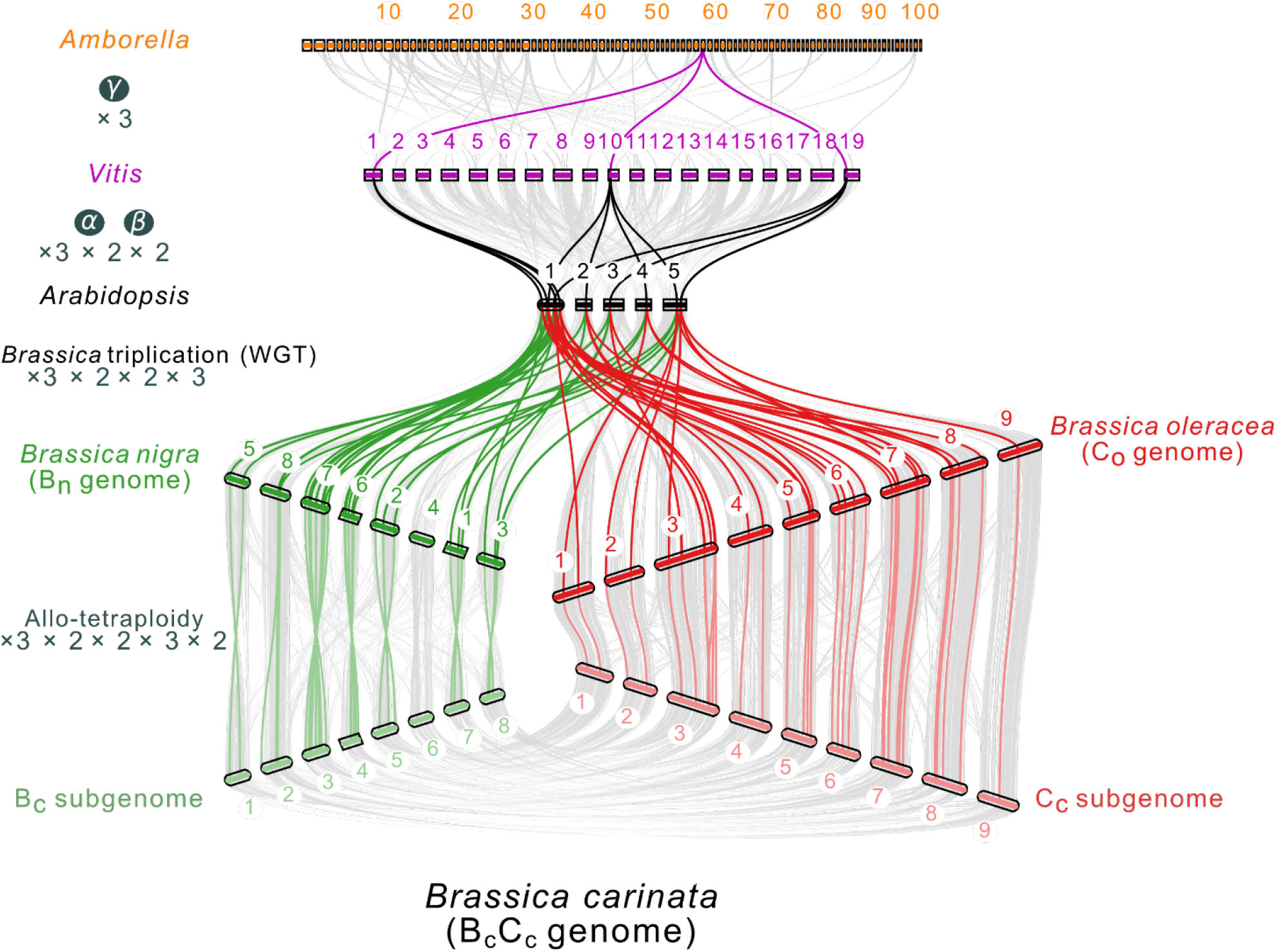
Graphic representation of local syntenic regions in support of whole genome duplication (WGD) events affecting *Brassica* species. Syntenic gene arrangement for the basal angiosperm, *Amborella trichopoda* (2n=100); the basal eudicot, *Vitis vinifera* (2n=19); and the model plant *Arabidopsis thaliana*, as well as the two progenitor species, *Brassica nigra*, *B. oleracea*, and *B. carinata*. Grey lines connecting the species indicate conserved syntenic blocks containing more than 30 orthologs. The highlighted lines represent the homologs among the genomes that originated from one *Amborella* gene after WGD events including the gamma WGD, the alpha and beta WGD, the *Brassica*-specific WGT (whole-genome triplication), and the *B. carinata*-specific hybridization between *B. nigra* and *B. oleracea*. The concept of this figure was adapted from Chalhoub, B. *et al* (2014).

We compared TE classes to identify any repetitive elements or trends specific to the *B. carinata* genome. LTR retrotransposons contributed the most considerable fraction of repetitive elements, covering 26.19% of the total genomic DNA of *B. carinata* (Table S8) *Copia* and *Gypsy* elements represented the major superfamilies, covering 8.8% and 14.7% of the genome, respectively (Table S10). An asymmetry in LTR content was observed with LTR sequences contributing to 14.1% and 12.1% of the Bc and Cc subgenomes, respectively (Table S10). An asymmetric LTR distribution was also identified between the An and Cn subgenomes of *B. napus* (Chalhoub et al., 2014; Parkin et al., 2014). The dominant Cn subgenome comprises a higher LTR proportion (8.55%) compared to the smaller, relatively gene-poor An subgenome (5.86%, Figure 3F). These results are consistent with previous analyzes of the repeat content of the progenitor genomes *B. nigra* and *B. oleracea* (Figure 3C) (Perumal et al., 2020; Luijten, 2010). Similar to the trends in gene content among the six species, the A (sub)genomes generally have a lower LTR content and the C (sub)genomes generally have a higher LTR content. While their B (sub)genomes show the greatest variety in LTR content, the majority of the B (sub)genome assemblies have the highest LTR content, with both *B. carinata* Bc subgenome assemblies having a higher LTR content with greater variation between the two assemblies and the Cc subgenome assemblies have a LTR content with less variation between them.

Notably, the *B. carinata* genome contains the lowest *Helitron* content among the *Triangle of U* B and C subgenomes. As *Helitron* content is generally lower and more stable than that of RNA transposons in plant genomes, *Helitron*-related data could be more informative of plant genome features (Hu et al., 2019). Although *Helitrons* compose 15.2% of related *Brassica* genomes (Bj, Cn, Bn, and Co), *Helitrons* only take up 8.2% of the *B. carinata* genome (Table S8-S10). While the *B. carinata* subgenomes contain a lower *Helitron* proportion (ANOVA, p>0.001), there is no statistical significance in the total abundance of *Helitrons* between *B. carinata* and its related (sub)genomes (ANOVA, p=0.66) or sum length in bp (ANOVA, p=0.88). This implies a disproportionate expansion of the *B. carinata* genome relative to *Helitron* content. *Helitrons* are thought to participate in loss of function and generation of diversity in coding regions due to insertion mutations and exon shuffling (Morgante et al., 2007; Gao et al., 2016). We hypothesize that the reduced *Helitron* content in the *B. carinata* genome might have facilitated higher levels of gene family expansion. Although the transposition mechanism of *Helitron* remains unclear (Brunner et al., 2005), the domestication of this crop could have benefitted from a limitation to this process, possibly allowing the higher levels of gene family expansion we identified.

The contiguity of our assembly was assessed in terms of intact LTR retrotransposons using the LTR Assembly Index (LAI), which uses the identification of intact LTRs as a proxy for assembly contiguity (Table S11). The Gomenzer assembly had an LAI score of 12.23 for the whole genome, with the Bc subgenome having an LAI of 11.78 and the Cc subgenome containing an LAI of 12.71. Intact LTR retrotransposons might have been primarily inherited from the diploid progenitor genomes. These findings guided us towards further investigation into biased repeat content and differences in intact LTR numbers between the (sub)genomes as possible explanations for observed genome dominance patterns (Hollister and Gaut, 2009; Edger et al., 2017; Bird et al., 2020).

### The Zd-1 *B. carinata* genome assembly lacks the same contiguity and completeness of our Gomenzer assembly

Another *B. carinata* genome assembly (accession Zd-1) was published in 2021 (Song et al., 2021). However, our intergenomic alignment a lack of collinearity between the two assemblies. The differences are particularly apparent in the Bc subgenome where there are truncated chromosomes (GomB01, GomB02, GomB05), large segmental inversions (GomB03, GomC03, GomC04), and a duplication of chromosome GomB06 (Figure S36). While some inconsistencies are expected due to true variation between the accessions, our assembly shows superior quality metrics and is likely a more complete reference genome. When comparing the two assemblies, only a small discrepancy in BUSCO scores is found at the whole genome level, the Zd-1 scores drop considerably when considered by subgenome (Figure S10). As the *B. carinata* genome is tetraploid, a high presence of duplicated BUSCOs is expected with this highly redundant genome. while the Zd-1 assembly contains 9% fewer BUSCOs, our Gomenzer assembly contains 49% more duplicated BUSCOs. The Zd-1 Bc subgenome contains 31% less BUSCOs than the Gomenzer Bc subgenome and the Zd-1 Cc subgenome contains 10% less BUSCOs.

Our 1.31 Gbp Gomenzer assembly was evaluated to be 20.52% larger than the 1.09 Gbp Zd-1 assembly. In particular, the Bc subgenome showed the largest genome size difference between Gomenzer and Zd-1. The difference in genome size is not proportionate, with the Bc subgenome lacking a significantly larger proportion (48.4%) compared to the Cc subgenomes (12.0%). Likewise, our Gomenzer assembly has a higher gene content with 22.8% more annotated genes, particularly in the Bc subgenome which contains 72.90% more genes identified than its Zd-1 counterpart (Table S13). Thus, both genome size and gene content differences between Gomenzer and Zd-1 were most pronounced in the Bc subgenome. Moreover, our Gomenzer assembly classifies as reference quality under the LAI classification system (10 ≤ 12.23 < 20), while the Zd-1 could be defined in the draft quality type (0 ≤ 9.57 < 10) through the LAI genome classification system (Ou et al., 2018). As such, our assembly Gomenzer assembly has higher continuity than the Zd-1 assembly in terms of repetitive sequence space. Overall, our *B. carinata* var. Gomenzer assembly genome size result is likely more accurate and reliable compared to the Zd-1 assembly. Specifically, Zd-1 assembly is missing large portions of sequence and exhibits genome structural disparity between respective progenitors. Considering the difference in genome contiguity and incompleteness, the Zd-1 assembly might include artifacts from the assembly process rather than true variation between the accessions.

### Our high-quality genome assembly refines estimations into the timing of *Triangle of U* speciation events

The *Triangle of U* genomes are highly duplicated, with diploids having undergone an aggregate 36⨉ multiplication (3 ⨉ 2 ⨉ 2 ⨉ 3) and the allotetraploids having undergone an aggregate 72⨉ multiplication (3 ⨉ 2 ⨉ 2 ⨉ 3 ⨉ 2) since the most recent common ancestor of all eudicots (Figure 5). A *Brassica-*specific whole-genome triplication (WGT) event is believed to be the root of the rich morphotype diversity in *Triangle of U* species, giving rise to the speciation of the diploid species *B. rapa* (Ar), *B. nigra* (Bn), and *B. oleracea* (Co). According to our fossil clock calibration with *Arabidopsis lyrata*, *Arabidopsis thaliana*, *Arabis alpina,* and *Raphanus sativus*, the Bc and Cc subgenomes likely appeared about 1.25–2.87 and 1.05–2.2 MYA from *B. nigra* and *B. oleracea*, respectively (Figure 6A). The divergence times of the other *Brassica* subgenomes, *B. juncea* (Aj and Bj) and *B. napus* (An and Cn), likely occurred about 1.2–2.06 and 1.22–1.67 MYA, respectively. These results imply that the subgenome progenitor of each allotetraploid *Brassica* might be different from that of modern diploid *Brassica* species. We deduced that the hybridization leading to speciation of *B. carinata* (BcCc) occurred about 11,000–29,900 years ago. The hybridization events leading to the speciation of *B. juncea* (AjBj) and *B. napus* (AnCn) occurred an estimated 11,000–28,600 and 5,600– 7,000 years ago, respectively (Figure 6D). These determinations are earlier than the previous estimates for *B. juncea* at 3,900–5,500 years ago (Wang et al., 2019; Yang et al., 2016), with speciation of *B. napus* showing a similar time frame (Chalhoub et al., 2014), corresponding to the self-comparison *K*s values. Major hybridization events provide insight into allotetraploid evolution, particularly because of the relationship between polyploidization and subsequent crop domestication and improvement, which is an active area of research (Chalhoub et al., 2014; Pires et al., 2004; Parkin et al., 1995; Udall et al., 2005; Trick et al., 2009; Paritosh et al., 2021; Sharpe et al., 1995).

**Figure 6.**
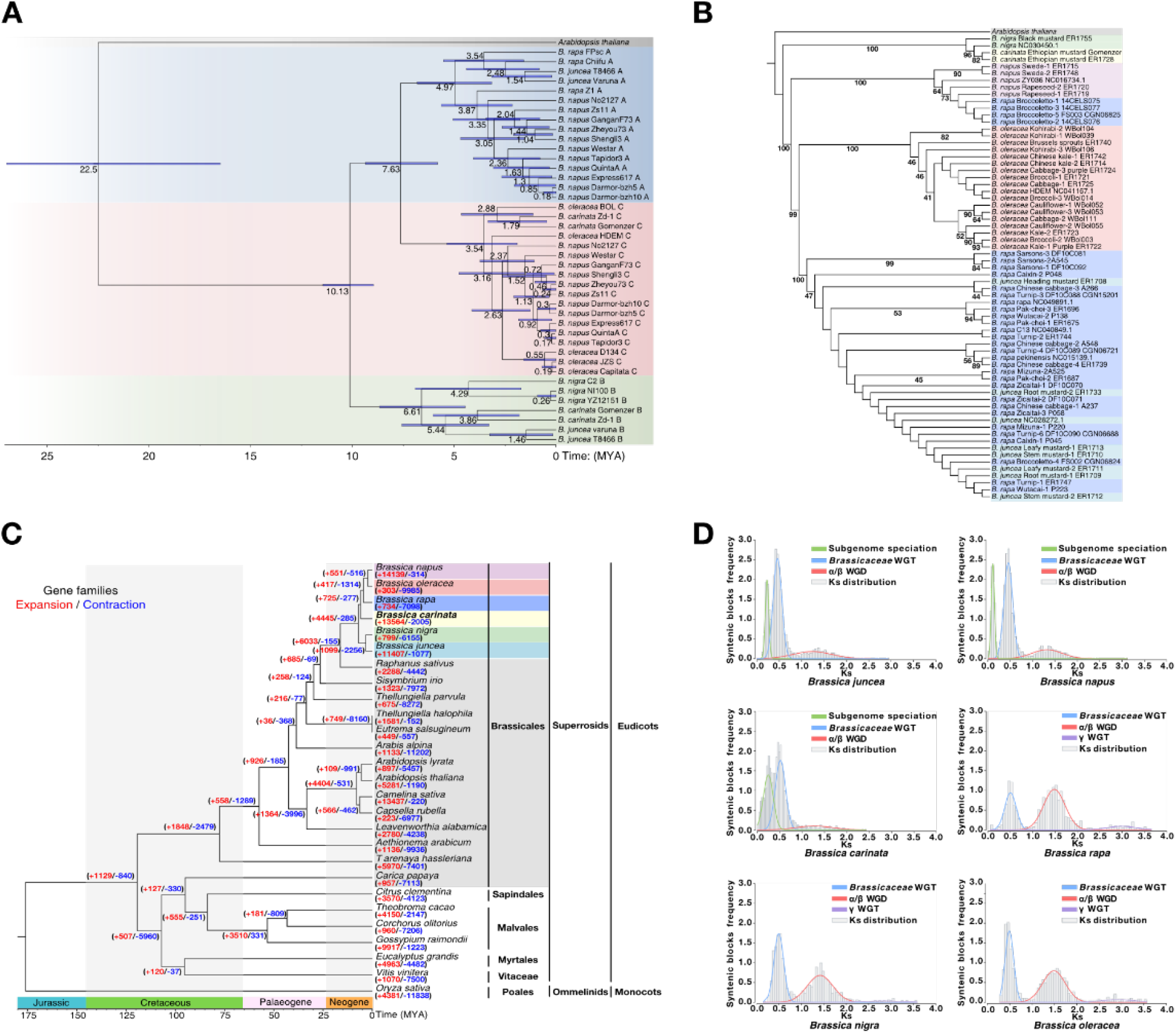
Phylogenetic trees among *Brassica* species and other dicot or monocot plant families. **(A)** The phylogenetic tree was constructed based on the orthologous low copy nuclear (LCN) genes between *Brassica* species with *Arabidopsis thaliana* as the outgroup. The species are highlighted in colors based on the three subgenomes (A, B and C). **(B)** A maximum likelihood (ML) phylogenetic tree of the *Brassica* chloroplast genomes with *Arabidopsis thaliana* as the outgroup. **(C)** A time-calibrated maximum likelihood phylogenetic tree among *Brassica spp.* and more distantly related species produced based on the expansion and contraction of orthologous genes. **(D)** Synonymous substitutions per site (*Ks*) values in *Brassica* species. The grey *Ks* distribution histograms represent *Brassica* synteny blocks. The γ whole-genome triplication (WGT) is indicated in purple, the red curve line indicates α/β whole-genome duplication (WGD) event from *Vitis* to *Arabidopsis*, the blue line represents *Brassicaceae* WGT where diploid *Brassica* species occurred from *Arabidopsis*, and the green line describes subgenome speciation event when the allotetraploid species were introduced.

To further elucidate the origin of the hybridized genome, we assembled the chloroplast genome from our genome sequencing data to identify the maternal progenitor of *B. carinata* (Figure S11). We used the concatenated coding sequence matrix to infer phylogenetic relationships. A distinct clade of only *B. nigra* (Bn) and *B. carinata* (BcCc) was identified, designating *B. nigra* as the maternal antecedent of *B. carinata,* according to maternal inheritance in *Brassica* (Figure 7B) (Scott and Wilkinson, 1999). The clear separation of *B. carinata* from other *Brassica* species implies considerable divergence of the chloroplast genomes belonging to the three diploid species. Both *B. napus* (AnCn) and *B. juncea* (AjBj) clustered with *B. rapa* (Ar) independently, supporting uniparental inheritance from *B. rapa* (Li et al., 2017; Kim et al., 2018; Xue et al., 2020). However, the phylogenetic placement of *B. oleracea* (Bo) between *B. napus* (AnCn) and *B. juncea* (AjBj) suggested some divergence in the maternal *B. rapa* parent (Ar) before the hybridization of the two allotetraploids. The different timing of the respective hybridization events that produced *B. napus* (AnCn) and *B. juncea* (AjBj) also supports maternal genome divergence. These observations are consistent with previous work suggesting that a turnip-type *B. rapa* (ssp*. rapa*) was the ancestor of *B. napus* (Lu et al., 2019) and a yellow sarson-type *B. rapa (*ssp. *trilocularis*) was the parent of *B. juncea* (Yang et al., 2016).

**Figure 7.**
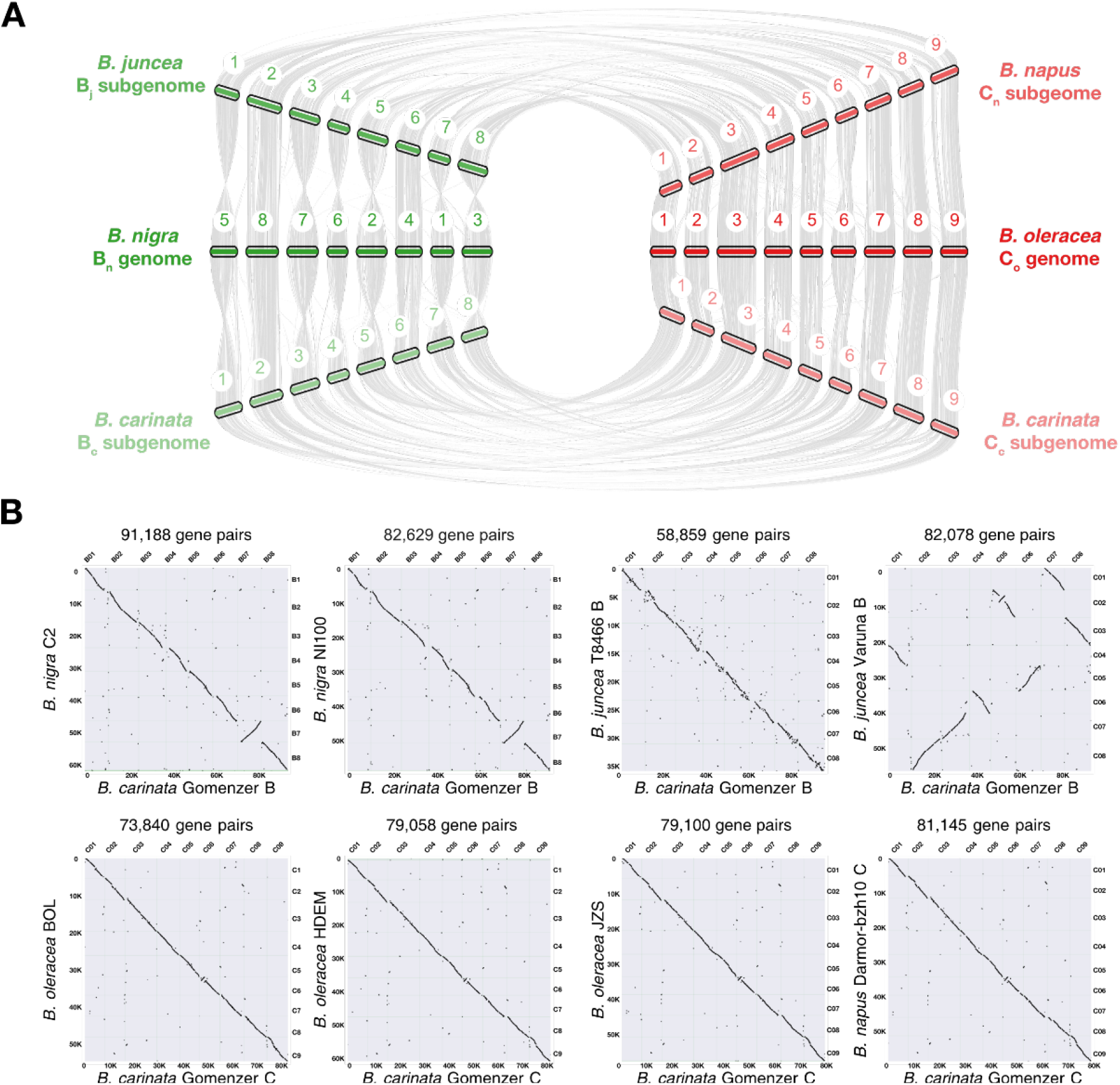
Genomic alignment between progenitor *Brassica* species (*B. rapa*, *B. nigra* and *B. oleracea*) and the hybridized allotetraploid species (*B. carinata*, *B. juncea* and *B. napus*). **(A)** Collinear relationship analysis at the subgenome level in *Brassica* species. The collinearity between respect ancestor *Brassica* species and respect subgenomes of allotetraploid species was analyzed based on the orthologous gene comparisons. **(B)** Pairwise intergenomic comparison dot plots between the two *B. carinata* subgenomes and their respective shared (sub)genome among other *Brassica* assemblies (Bn, Co, Bj, and Cn). The black dots indicate the syntenic genes conserved as a block among the two genomes. Longer lines represent a high abundance of genomic collinearity.

### Transcription factor gene family expansion facilitated gene neo- and subfunctionalization during *B. carinata* domestication

The *B. carinata* genome has undergone extensive gene family expansions (Figure 7C). Consistent with these observed gene expansions, moderate gene number gains and low gene number losses were identified in *Brassica* lineages across the species complex. Specifically, *B. carinata* showed an average gene expansion/gene family ratio of 3.16, reflecting the expansion of 5,301 gene families in its genome (Table S18).

While *B. napus* displayed the highest number of expanded gene families among the *Triangle of U* (7,153), *B. carinata* showed the highest gene expansion/gene family ratio (3.16 vs. 2.61). Pathway analysis of expanded gene families revealed enrichment of genes in signal transduction and environmental adaptation-related pathways. Notably, enrichments were observed in abiotic and biotic stress-related pathways, photosynthesis, plant hormone signal transduction, diterpenoid biosynthesis, plant−pathogen interaction, ubiquitin-mediated proteolysis, and ascorbate and aldarate metabolism (Yanagawa and Komatsu, 2012; Mathur et al., 2014; Wang et al., 2015; Mafu et al., 2018; Smirnoff, 2018). The expansion of these pathways suggests that *B. carinata* has developed effective adaptive responses to biotic and abiotic stresses (Figure S12). Gene Ontology (GO) analysis also revealed significant gene number enrichments related to gene regulation and transcription (Figure S14-S16). The retention of successive duplicated genes in multiple pathways might have contributed to the improved environmental adaptability and novel trait development observed in *B. carinata*.

A total of 8,653 transcription factors (TFs) across 81 TF families were identified in *B. carinata,* representing an expansion in the allotetraploid genome compared to its diploid progenitor genomes (Table S19, Figure S17). *B. carinata* exhibited the highest number of TFs among *Brassica* species in this study (8,653) followed by *B. napus* (7,342) (χ^2^, *P* = 2.35 x 10^-15^). Not surprisingly, more TF-encoding genes were found in the gene-rich Bc subgenome with 59 TF families possessing more genes in the Bc subgenome (4,757) than in the Cc subgenome (3,527) (χ^2^, *P* < 3.52 x 10^-8^). Similar TF gene family expansion and contraction patterns were evident among the allotetraploids, but not among the *Triangle of U* diploids (Table S18-S19) (Chalhoub et al., 2014). Such TF expansion implies an expansion of regulatory networks and, hence, the functional divergence of genes and regulatory networks during domestication. which might confer greater potential for environmental adaptability in the allotetraploids relative to their progenitor species.

Polyploidization triggers genome fractionation to compensate for the evolutionarily instantaneous duplication of all genes and cis regulatory elements (Zhang et al.). Gene fractionation, or the process through which homoeologous subgenomes reciprocally lose genes such that genes return to single-copy status, was also examined also in *B. carinata* (Renny-Byfield et al., 2017). A total of 6,887 and 4,761 genes from *B. napus* (Bn) and *B. oleracea* (Co), respectively, had no counterpart in *B. carinata* (Table S30). These numbers indicated relatively little gene loss compared to that experienced by shared-genome species, particularly from the B subgenome (Bj and Cn, 9,708 and 6,963, respectively). Several reports have emphasized that most lineages have undergone extreme gene loss or genome size reduction (BENNETT and LEITCH, 2005; Leitch and Leitch, 2008) and diploidization at many loci (Wolfe, 2001) following polyploidization. Fractionation rates are generally high in newly formed polyploids then decelerate over the next millions of years as the genome returns to a diploid state (Zhang et al.). Although *B. carinata* has undergone diploidization to some extent, a simple whole-genome comparison did not indicate substantial gene loss. With our estimation of hybridization at 11,000–29,900 years ago, *B. carinata* is a relatively new polyploid, or mesopolyploid, and only a small degree of gene fractionation has occurred compared. Interestingly, the *B. carinata* Bc genome has experience considerably less gene loss compared to despite our findings that its hybridization likely occurred prior to the other allotetraploids.

### Redundant gene evolution and selection pressures on duplicated genes among ***Brassica* genomes**

Duplicated genes can also diversify and take on new functional roles (Tang et al., 2010; Wu and Qi, 2010), and might also serve as substrates for increased adaptation to stressful environments (Alix et al., 2017). The majority of orthologous genes survived as homoeologous pairs in the *B. carinata* genome. A total of 84.6% of the total genes were identified as orthologous to their respective progenitors (Bn and Co) and shared-genome species (Bj and Cn, Table S25) A total of 8,315 collinearity blocks, or a run of syntenic orthologs between the compared genomes, were identified. Of these, 75,483 *B. carinata* genes could be aligned with 59,257 genes of the diploid progenitors (Bn and Co) while still maintaining synteny (Figure 7, Table S26). We found that the Bc subgenome had the highest proportion of homoeologs relative to the total gene content among the *Triangle of U* B genomes at with 65.4% of the Bc genes being homoeologs (Bn 46.3%, Bj 41.4%, Table 27). The Cc subgenome also had the highest homoeolog proportion among the C (sub)genomes (Cc 59.1%, Co 46.8%, Cn 40.2%). These results imply that the *B. carinata* genome, and particularly the Bc subgenome, is more prone to homoeolog retention than the other genomes.

We compared sequence evolution rates among homoeologous pairs by assessing the ratio of nonsynonymous substitutions (*Ka*) and synonymous substitutions (*Ks*), or the ω values (*Ka*/*Ks,* dN/dS). As expected, the homoeologs generally have larger *Ka* and *Ks* values in the *Brassica* diploid genomes relative to the tetraploids due to the increased amount of time since their emergence (Table S28). The tetraploid homoeolog ω values are larger than the diploid homoeolog ω values, indicating a greater degree of sequence divergence between homoeolog pairs in the allotetraploids (ANOVA, p>0.001). A greater degree of sequence evolution in the *B. carinata* genome compared the other allotetraploids. The *B. carinata* genome had a larger homoeolog ω value (0.31), than the other allotetraploids, with *B. napus* (AnCn) and *B. juncea* (AjBj) having homoeolog ω values of 0.28 and 0.26, respectively. The increased rate of divergence in sequence between *B. carinata* homoeologs relative to the other tetraploids suggests a relatively higher level of adaptive evolution, possibly facilitating the species’ high resilience to environmental stresses. Interestingly, the differences in ω values among the allotetraploids were more affected by differences in background evolution rates than differences in nonsynonymous mutations. For example, *B. juncea* has a relatively higher homoeolog *Ks* value, leading to the lowest ω value among the allotetraploids. The high background sequence evolution rate in *B. juncea* supports our finding that *B. juncea* emergence as a species earlier than previously thought.

Paralogs, or duplicated genes derived from means other than WGD events, were also assessed in the *B. carinata* genome. A total of 2,129 tandem, 8,508 proximal, 11,925 transposed, and 7,374 dispersed paralogs were identified (Table S27). A smaller proportion of tandem duplicates (TDs), which are linearly adjacent to each other, relative to the total gene content was identified in *B. carinata* compared to the related (sub)genomes. The Bc and Cc subgenomes contain a 2.4- and 2.1-fold lower TD content than their related (sub)genomes, respectively, indicating lower rates of TD propagation and/or retention (Table 27). Interestingly, the most extended TD array was located in the Cc subgenome. For all six species, the TD ω values are about 1.8-fold larger than the WGD ω values, indicating a higher diversifying positive selection on TDs. Although *B. napus* (AnCn) has the highest TD ω value (0.53) among the *Triangle of U*, the *B. carinata* TD ω value (0.39) is still larger than those of the respective progenitor genomes (0.33). One of the most overrepresented GO terms among the TDs was ‘DNA methylation on cytosine within a CG sequence’ (Figure S28), which is associated with epigenetic modification and stress adaptation through reprogramming the transcriptome and altering genome stability that could enhance resilience to stress (Tirnaz and Batley, 2019). Thus, the *B. carinata* TDs suggest that frequent functional diversification (neofunctionalization and/or subfunctionalization) has occurred in this species after genome hybridization. The functional divergence of the relatively small number of TDs might have helped *B. carinata* adapt to changes in its environment.

We identified 4-fold more proximal duplicates (PDs), which are separated by a few genes, than TDs. Likewise, a larger proportion of PDs relative to the total gene content was identified in *B. carinata* compared to the respective progenitor genomes (Bn and Co) and shared-subgenome species (Bj and Cn, Table 27). The Bc and Cc subgenomes contain a 2.0- and 1.4-fold higher PD content than their related (sub)genomes, respectively. We found a similar sequences evolution rate for PDs as the TDs, concurrent with the current literature (Qiao et al., 2019). However, we found that PD sequences diversify more rapidly in *B. carinata* (0.52 ω value) compared to the other five *Triangle of U* species (Table S28). The higher PD ω value appears to be a result of an unusually low *Ks* value for *B. carinata* PDs, with a mean *Ks* value of 0.30 compared to the average *Ks* value of 0.52 for all six species. These results suggest that the *B. carinata* genome contains a relatively high number of young PDs that might have occurred relatively more recently in *B. carinata,* yet diversify quicker than the PDs of related genomes. The GO terms enriched among PDs were associated with lipid metabolism and interspecies interactions, including those with viral pathogens (Figure S27). The ω values of the more distant transposed duplicates and dispersed duplicates lay at the median of the five duplication modes considered. However, the *Ks* values of these distant duplications were up to 3.5-fold greater than the homoeolog *Ks* values, indicating a longer period of time has passed since their duplication.

### Genes encoding key erucic acid synthesis**-**related enzymes display Bc subgenome dominant expression

We then investigated agronomically important genes targeted in *Brassica* improvement efforts to test whether the dominant Bc subgenome shows higher expression as the subgenome dominance hypothesis describes (Freeling et al., 2012). With respect to acyl lipid metabolism of this oilseed crop, genes in the fatty acid biosynthesis and elongation pathways have undergone notable expansions in *B. carinata* (Table S21). In higher plants, *de novo* fatty acid synthesis begins with the conversion of glycolysis-derived pyruvate from acetyl-CoA by the chloroplast-localized acetyl-CoA carboxylase (ACCase) complex. ACCase is the rate-limiting step in fatty acid (FA) synthesis and has been regarded as a major bottleneck in triacylglycerol (TAG) biosynthesis, the major form of lipid storage in the seeds (Ke et al., 2000). The three core enzymes of the ACCase complex have been reported to maintain a constant stoichiometric ratio of their transcript abundance, which was observed in the silique transcriptome data (*BCCP1*/*CAC1*, 71.7 TPM; *CAC2*, 68.5 TPM, *CAC3*, 66.2 TPM, Figure 8). Still, we identified Bc subgenome biased expression of the three nuclear-encoded genes for the ACCase complex in developing siliques, contributing 50.4% - 58.3% of the total transcripts (Ke et al., 2000).

**Figure 8.**
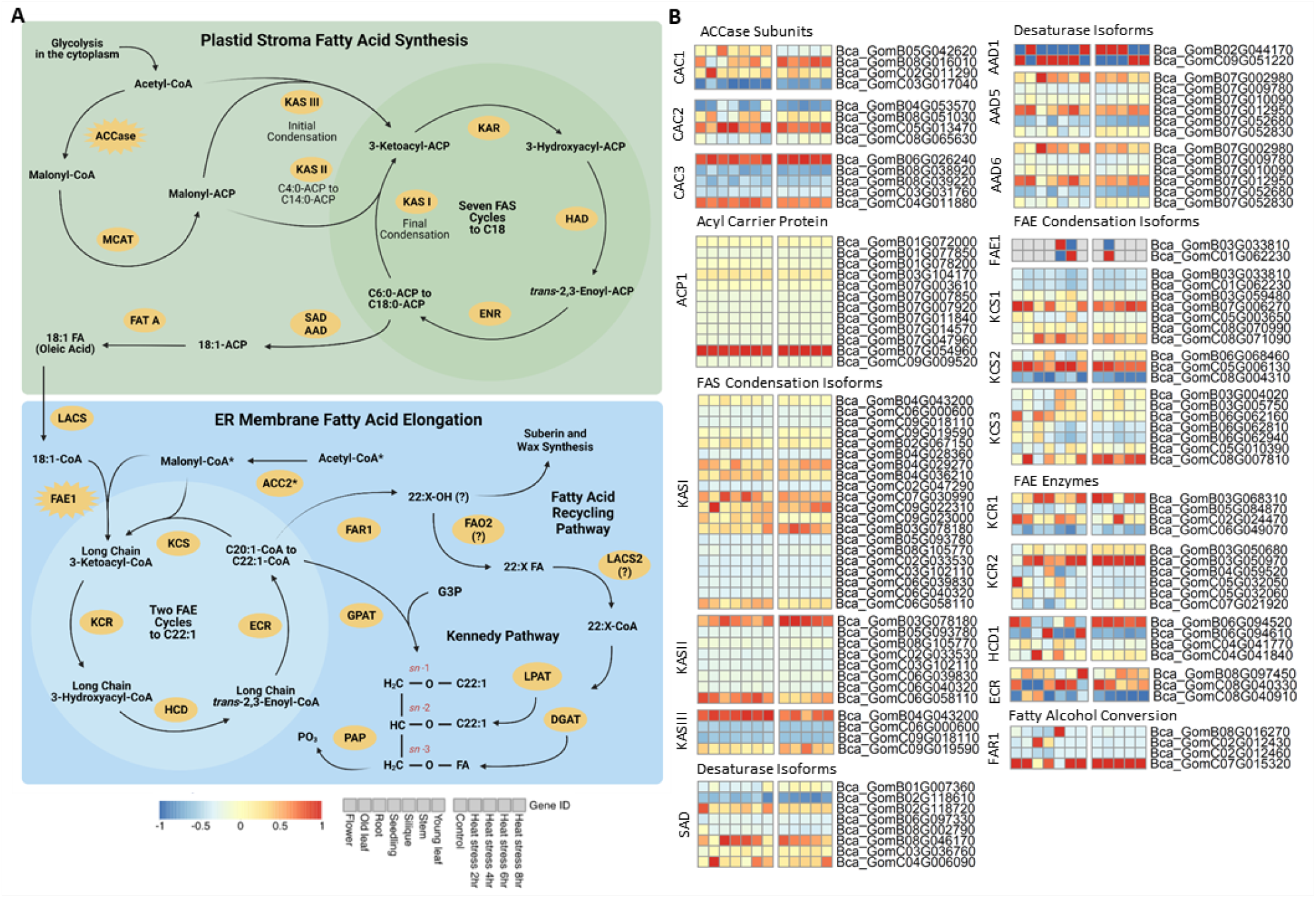
Erucic acid triacylglycerol biosynthesis pathway. **(A)** Fatty acid synthesis, elongation, and triacylglycerol assembly, and fatty acid recycling. **(B)** Relative expression level of genes involved in erucic acid biosynthesis. The relative abundance of gene expression is visualized as a heat map, ranging from blue to red with increasing relative expression levels. The x-axes of the heat maps describe the tissue or heat stress time point from which the RNA was extracted and the y-axis provides the gene IDs. Gene abbreviations: PDHC, pyruvate dehydrogenase complex; ACCase, Acetyl-CoA carboxylase complex; ACP, acyl carrier protein; MCAT, Malonyl-CoA ACP S-malonyltransferase; KAS, β-ketoacyl-ACP synthase; FAS, fatty acid synthesis; KAR, β-ketoacyl-ACP reductase; MTHD, mitochondrial β-hydroxyacyl-ACP dehydrase; ENR, enoyl-ACP reductase; SAD, stearoyl-ACP Δ9-desaturase; FAT A, fatty acyl-ACP thioesterase A; LACS, long-chain acyl-CoA synthetase; KCS, β-ketoacyl-CoA synthase; KCR, β-ketoacyl-CoA reductase; HCD, β-hydroxyacyl-CoA dehydrase; ECR, enoyl-CoA reductase; *FAE*, fatty acid elongation; G3P, glycerol-3-phosphate; GPAT, glycerol phosphate acyltransferase; LPA, lysophosphatidic acid; LPAT, lysophosphatidyl acyltransferase; PA, phosphatidic acid; PAP, phosphatidic acid phosphohydrolase; DAG, diacylglycerol; DGAT, diacylglycerol acyltransferase; TAG, triacylgycerol.

The resulting acetyl-CoA is converted to malonyl-ACP, with the acyl carrier protein (ACP) acting as a shuttle for the acyl intermediate and is essential to their elongation by the cyclical fatty acid synthase complex (FAS) [119]. ACP is encoded by *ACP1*, which is significantly expanded in the larger, more gene-rich Bc subgenome (11 homologs). Many of the Bc subgenome *ACP1* homologs are tandem repeats with three copies on chromosome B01 and seven copies on chromosome B07. *ACP1* is ubiquitously expressed across all tissues with one Bc subgenome copy (Bca_GomB07g054960) showing the highest transcript levels.

Beginning with an initial acetyl-CoA, each cycle of the fatty acid synthase (FAS) complex uses malonyl-ACP to add two carbons to the rowing acyl chain sequentially. The complex is composed of four proteins performing sequential reactions. The initial condensing enzyme, *KASIII,* and the genes encoding the next sequential enzymes of the FAS complex also displayed Bc subgenome biased expression in the developing siliques, with the Bc subgenome contributing 52.3% - 100% of the total transcripts. However, the isoforms that facilitate the subsequent FAS condensation reaction (*KASI* and *KASII*) showed Cc subgenome biased expression. Seven FAS cycles produce stearic acid (C18:0), which is subsequently desaturated before export to the cytosol. Four Δ9 stearoyl-ACP desaturases (SADs) that produce oleoyl-ACP, which work redundantly during seed oil storage, are found in the Arabidopsis genome. These SADs play a role in determining the resulting seed oil fatty acid composition and total concentration in Arabidopsis (Kazaz et al., 2020). The *B. carinata* homologs of these four SADs (*FAB2*, *AAD1*, *AAD5*, and *AAD6*) are significantly expanded in the Bc genome (13 copies) compared to the Cc subgenome (3 copies). Three of the four isozymes (*FAB2*, *AAD5*, *AAD6*) exhibit expression tissue-specific dominance of the Bc subgenome copies, with the Bc subgenome contributing 89% of the total transcripts for these genes in developing siliques and throughout eight hours of heat stress.

### FAE1 is a regulator of erucic acid synthesis

After export to the cytosol, oleic acid (C18:1 ^Δ9^)-CoA encounters the endoplasmic reticulum (ER) membrane-localized fatty acid elongation (FAE) complex, which extends FAS-derived FAs into very long-chain fatty acids (VLCFAs), such as erucic acid (C22:1 ^Δ13^), from the chloroplast-derived FA pool. The FAE complex is comprised of four membrane-bound proteins that perform sequential reactions, with each cycle adding two carbons to the growing acyl-CoA chain. The first of these four enzymes, ß-ketoacyl-CoA synthase (KCS), facilitates the condensation of a long to very-long chain acyl-CoA with a malonyl-CoA and is the rate-limiting step in VLCFA production. The resulting VLCFA species carbon chain lengths are determined by the substrate specificity of the numerous KCS isoforms (Millar and Kunst, 1997). *FAE1* is the isoform that targets oleic acid for elongation and governs erucic acid production in *Brassica* species (Saini et al., 2019). The *B. carinata* genome has retained only one *FAE1* homolog per subgenome. The other KCS-encoding genes (*KCS1-3*), which produce VLCFAs other than erucic acid (Guo et al., 2016), are relatively more expanded, indicating purifying selection pressure on *FAE1*.In contrast to the substrate-specific nature of KCSs, the other three the FAE complex enzymes (KCR, HCD, ECR) participate in the elongation of all VLCFAs (Millar and Kunst, 1997). However, the genes encoding these enzymes have multiple copies in the *B. carinata* genome that display tissue-specific expression, implying possible subfunctionalization (Figure 8).

*FAE1* expression is restricted to the developing siliques, stem, and early heat stress (2 hour) in *B. carinata.* The Bc subgenome copy, Bca_GomB03g033810, contributes 57.4% of the silique transcripts whereas the Cc subgenome copy, Bca_GomC01g062230, contributes 75% of the transcripts in the stem and is heat stress-responsive (Table S40). The silique-dominant Bc copy has a Tc1/mariner insertion in the promoter region and the stem-dominant Cc subgenome copy has two PIF/Harbinger insertions. Whereas Gomenzer is a high erucic acid accession of *B. carinta,* PIF/Harbinger-like DNA transposon insertions in the *FAE1* promoter are correlated with low erucic acid accumulation in *Sinipis alba* (Zeng and Cheng, 2014). These results suggest that the dominant Bc subgenome contributes more to seed oil biosynthesis whereas the Cc subgenome might be more involved in modulating leaf membrane lipid composition in response to heat stress and stem membrane lipid composition under normal development (Zhukov and Shumskaya, 2020; Zoong Lwe et al., 2021). As *FAE1* expression is generally limited to developing seeds (Zeng and Cheng, 2014), *FAE1* activity in the stems and in response to heat stress may contribute to *B. carinata*’s high heat and drought tolerance.

### *FAE1* homologs among the A, B, and C (sub)genomes

Although *FAE1* polymorphisms are known to have a key role in the level of erucic acid production in *Brassica* species (Millar and Kunst, 1997), there is debate about whether CDS or promoter region polymorphisms are the predominant mechanism underlying the high or low erucic acid phenotypes due to contradictory evidence from different *Brassicas spp*. (Saini et al., 2019). However, the increase in quality of plant genome assemblies in recent years has improved our ability to assess PAVs, in contrast to SNP analysis which has been extensively implemented in genetic studies without the aid of single-molecule sequencing technologies. The putative *FAE1* promoter varied from 1.0 to 1.3 kbp upstream of the transcription start site among the *Brassica* diploids in previous reports (Saini et al., 2019), which warranted PAV analysis on this region.

The *FAE1* homologs tended to situate themselves similarly among the *Triangle of U* genomes with the majority of the A, B, and C (sub)genome homologs placed on chromosomes A08, B03, and C03, respectively, suggesting conservation of copy number and locus. Each A (sub)genome homolog is positioned within an eight Mbp region on chromosome A08 with no duplicates. The *B. rapa* (Ar) Z1 assembly is the only A (subgenome) homolog lacking a promoter TE insertion. Similar to the A (sub)genome homologs, all of the C (sub)genome homologs are located on a 19 Mbp region of chromosome C03, except for those of *B. carinata,* which were transferred to chromosome C01. While upstream *Helitrons* were the prevailing insertions among the three (sub)genomes, the A (sub)genomes were enriched for hAT insertions and the C (sub)genomes were enriched for PIF/Harbinger insertions, both of which are classified as MITEs. Eight of the ten *B. napus* (AnCn) genomes investigated shared a 3,074 bp hAT insertion in the promoter region of the A08 homolog, including No2127, the only high erucic acid accession. Considering this hAT insertion is present in the high and low-erucic acid accessions, the No2127 C subgenome copy likely facilitates erucic acid production and is engulfed by a *Helitron*, possibly carrying a promoter or enhancer sequence.

The *B. juncea* (AjBj) vegetatable T844 accession contains two *FAE1* homologs: one on chromosome B03 and the other on chromosome B04, both of which contain a *Helitron* insertion. We also confirmed the two *FAE1* homologs in the oleiferous Varuna accession (Gupta et al., 2004): one on chromosome A08 preceded by a *Helitron* insertion and the other on chromosome B07 with no TE insertions in the promoter region. Deletion of an AT-rich region in A (sub)genome *FAE1* homologs is correlated with low erucic acid in *B. rapa* (Ar) and *B. juncea* (AjBj) (Saini et al., 2019). *Conversely, Helitrons*, which display an insertion preference for AT-rich regions (Grabundzija et al., 2016), can function as enhancer elements in tissue-specific promoters of higher plants (Sandhu et al., 1998) such as *FAE1,* whose expression is generally restricted to developing seeds (Zeng and Cheng, 2014). Unlike T8466, Varuna is a high erucic acid accession (∼47%) (Gupta et al., 2004) which might be facilitated by the lack of a *Helitron* insertion in the promoter region of the single *FAE1* copy in the dominant Bj subgenome, freeing the gene from the *Helitron-*mediated repression.

### A fatty alcohol diversion pathway for erucic acid biosynthesis as a possible

Two FAE cycles produce erucoyl-CoA (22:1^Δ13^) from oleoyl-CoA (18:1^Δ9^) in the ER membrane. Erucoyl-CoA is transferred to a glycerol-3-phosphate (G3P) backbone to form seed storage triacylglycerols (TAGs). After the addition of two acyl-CoA molecules to the glycerol backbone, the resulting TAG is packaged into an oil body that is pinched off from the ER membrane. Erucic acid is typically encountered on the *sn-1,3* positions of TAG but can occupy the *sn-2* position as well in the case of trierucin, a TAG with erucic acid at all three positions on the glycerol molecule (Lassner et al., 1995). Alternatively, VLCFA acyl-CoAs can be converted to fatty alcohols by ER-localized fatty acyl-CoA reductases (FARs) which can be directed to suberin or cuticular wax in their free form or, in the case of cuticular waxes, as wax esters [104]. FARs have unique acyl chain lengths and chain saturation specificities. *AtFAR1* primarily converts C22:0 fatty acyl-CoAs to C22:0 fatty alcohols [104], [105]. However, the fatty alcohols produced by FARs are also dependent on the composition of the available fatty acyl-CoA pool, making the resulting fatty alcohol composition dependent upon the species in which they are expressed [106]. For example, the jojoba FAR produces erucoyl alcohol (C22:1-OH) when expressed in a high erucic acid *B. napus* accession, diverting a portion of the fatty acids from TAGs to wax esters (Metz et al., 2000). In *B. carinata,* erucoyl-CoAs are likely converted to erucyl alcohol by *FAR1*.

The function for *FAR1* in erucic acid biosynthesis is not intuitive as *FAE1.* Apart from the jojoba FAR involved in the synthesis of fatty alcohol precursors for wax ester synthesis, a storage lipid vastly different from the TAGs found in Brassica and many other species, plant FARs are mainly known for their function in generating fatty alcohols destined for cuticular waxes, suberin, and pollen exine (Rowland and Domergue, 2012). One plausible route for *FAR1* contribution to seed erucic acid biosynthesis is through a fatty alcohol diversion pathway in which a portion of erucoyl-CoA is reduced by *FAR1.* The resulting erucoyl alcohol is subsequently oxidized to erucic acid, perhaps through an aldehyde intermediate, which can then be reincorporated into TAG after activation to erucoyl-CoA by a long chain acyl CoA synthetase (LACS) enzyme (Figure 8). Candidates for the oxidation of erucoyl alcohol to erucic acid and the reactivation of erucic acid to erucoyl-CoA include long-chain alcohol oxidases (FAO) and LACS, respectively, which have been identified in Arabidopsis, jojoba, and other species (Rajangam et al., 2013; Zhao et al., 2008; Cheng et al., 2004; Zhao et al., 2021).

The *B. carinata* genome contains four *FAR1* homologs: one copy in the Bc subgenome and three copies in Cc subgenome (Table S42). The Bc subgenome homolog (Bca_GomB08G016270) showed the highest expression, contributing 84% of the total transcripts across all tissues (Figure 8). The expression of this homolog is confined to the developing siliques with 79.9 TPM, which could indicate a possible function in embryo cuticle biosynthesis or even oil synthesis. All four of the *FAR1* homologs have multiple TE insertions in the promoter region, but the highly expressed copies contain CDS TE insertions. Unlike the other *FAR1* homologs, Bca_GomB08G016270 contains multiple TE insertions in its CDS, including *Helitrons,* PIF/Harbingers, and a Mutator element (Table S42). Two of the three Cc subgenome homologs show low levels of expression with the most active copy, Bca_GomC07G015320, still only contributing 12.9% of the total transcripts across all tissues. Bca_GomC07G015320 contains a hAT insertion in the CDS while the less active. In contrast to our findings, *Helitron* insertions in *FAR1* genes have been associated with low erucic acid content in specific cultivars of *B. napus* (Hu et al., 2019).

### Gene family expansion and transposable element signatures in *FLC* homologs facilitate adaptation to different climates for crop expansion

Of the allotetraploids, *B. carinata* and *B. juncea* are both winter crops (Paritosh et al., 2021), while *B. napus* originated as a winter crop from which spring and semi-winter ecotypes have been developed through traditional breeding methods (Song et al., 2020). Late-flowering (winter) ecotypes overwinter prior to flowering to coordinate their reproductive development with the spring season. This process of cold exposure-induced flowering, or vernalization requirement, is facilitated in part by strong *FLOWERING LOCUS C (FLC)* expression. *FLC* is a MADS-box TF that is a key repressor of flowering that acts in a dosage-dependent manner and is one of the most important genes controlling the initiation of flowering in *Brassica* (Song et al., 2020).

Winter ecotypes typically display strong *FLC* expression, and therefore a strong vernalization requirement, while spring types flower earlier in the growing season due to weak *FLC* expression. A weak vernalization requirement allows adaptation to climates prone to summer drought, facilitated in part by SVs in *FLC,* a gene that is particularly prone to TE insertions (Quadrana, 2020).

TE insertions in the *FLC* promoter region and CDS can either upregulate [107] or downregulate [108] its expression. Song *et al*. (2020) found that presence or absence variations (PAVs) in one of the nine *B. napus FLC* copies (*BnaA10.FLC*) helped classify accessions by ecotype, a crucial aspect of crop breeding (Song et al., 2020). Eight copies of the *FLC* gene family were present in the *B. carinata* genome, which is expanded compared to the two and three copies found in *B. nigra* (Bn) and *B. oleracea* (Co) (Figure S19). The Bc subgenome contains five copies, however, none are expressed except for Bca_GomB08g003570, which contains three CACTA insertions within 3 kb of the TSS (Table S41). Two of the three Cc subgenome copies are expressed with Bca_GomC09g049840 contributing 77.2% of the *FLC* transcripts across all tissues. Overall, the Cc subgenome contributes 12.5-fold more FLC transcripts than the Bc subgenome. Interestingly, our phylogenetic analysis of the *Triangle of U FLC* homologs revealed that the most highly expressed copy in the *B. carinata* genome, Bca_GomC09g049840, resides in the same clade as *BnaA10.FLC* (GSBRNA2T00135921001) along with two other A (sub)genome homologs from *B. rapa* (Ar) and *B. juncea* (AjBj) (Figure S19). Both of the expressed Cc subgenome copies have an enrichment of Helitron insertions in the promoter region and the most highly expressed copy also contains a Gypsy retrotransposon insertion spanning the last 93 bp of the CDS (Table S41).

CACTA and *Helitron* TEs are miniature inverted-repeat transposable elements (MITEs) are non-autonomous, relatively short, A/T-rich DNA transposons. MITES are preferentially inserted in genes and gene flanking regions in plant genomes, leading to the speculation that these TEs might play an important role in genome evolution (Liu et al., 2019; Zerjal et al., 2009). Although they generally downregulate the expression of proximal genes likely due to TE hypermethylation or MITE-derived small RNAs, instances of MITE-mediated individual gene upregulation have been reported. Song *et al*. (2020) found that a MITE insertion in the promoter region of *Bna*A10.*FLC* was correlated with increased transcript abundance, possibly facilitating the development of winter ecotypes. Therefore, the MITE insertions in the promoter regions of the active FLC copies in *B. carinata* might promote their transcription, strengthening the vernalization requirement of this winter crop.

Interestingly, the submissive An subgenome in *B. napus* (AnCn) has a stronger influence on flowering time (Song et al., 2020), reflecting our finding of *FLC* expression dominance of the submissive Cc subgenome in *B. carinata*. This phenomenon could be attributed to the relaxed purifying selection pressure observed in the submissive subgenomes of allopolyploid species. The decreased selection pressure allows the retention of structural variants (SVs) such as TE insertions and gene copy number variations, which are known to impact phenotype to a greater degree than SNPs.

Moreover, the submissive subgenomes tend to have a higher global TE content due to increased TE retention (Alger and Edger, 2020), where TE insertions causing favorable mutations are more likely to be retained. Alternatively, subgenome dominance can be spatiotemporal-specific, with each subgenome facilitating discrete tissues or developmental stages despite global subgenome dominance patterns (Colle et al., 2019; Flagel et al., 2008). The overall submissive An and Cc subgenomes of *B. napus* and *B. carinata*, respectively, might contribute more to flowering time.

## Discussion

*B. carinata* displays great potential as a climate-resilient crop for use in semi-arid and tropical sub-humid environments. Patterns of gene expansion and functional divergence were characterized that might have spurred genetic novelty provided the basis for improvement of adaptive traits in this drought- and heat-tolerant species. Gene duplication has long been known to contribute to the genesis of novel traits, phenotypic variation, and adaptation to changes in the environment (Rizzon et al., 2006; Ohno, 1970; Kliebenstein, 2008). The *B. carinata* genome displays extensive gene duplications in families and pathways associated with important agronomic traits, including adaptation to semi-arid climates, resistance to diseases and pests, and synthesis of secondary metabolites with human health-promoting effects. For example, genetic pathways involved in the biosynthesis of cuticular waxes have undergone notable expansions in *B. carinata*. The accumulation of thick cuticular waxes, along with other key traits shared by *B. carinata*, are critically important for drought- and heat-tolerant species. These genes could serve as potential targets for future engineering and breeding efforts to improve climate resilience in other crops, especially the other species that are part of the *Triangle of U* model (Wang et al., 2020).

Unprecedented retention of transposon insertions in gene regions also generally follows polyploidization (Baduel et al., 2019), introducing, in part, the intraspecific diversity required for crop domestication. Structural variants (SVs), such as TE insertions, are major drivers of phenotypic variation. Therefore, TE insertions and copy number variations (CNVs) were analyzed in gene regions controlling key agronomic traits among the *Brassica* subgenomes. Transposon inserted into exon genes can alter gene structure and function by promoting chromosomal translocation and exon shuffling (Mobile Elements: Drivers of Genome Evolution), suggesting protein abundance of *Brassica* gene expression related to the initiation of flowering timing and acyl metabolism might be affected by TEs insertion. However, more detailed information about the effect of TEs on gene expression patterns is needed and is the subject of ongoing research.

Polyploidization events relax the purifying selection placed upon the newly merged genomes, often in a subgenome-specific manner with the submissive subgenome undergoing a higher relaxation of selective constraints (Bird et al., 2018). Subgenome dominance occurs in interspecific hybrids where there is biased expression among divergent genomes contained in one nucleus, leading to biased fractionation of the lowly expressed subgenome (Cheng et al., 2018). Rapid genomic downsizing is unique to angiosperms and might be facilitated through subgenome dominance (Cheng et al., 2018). Thus, some speculate that subgenome dominance appears to be angiosperm-specific might be at the core of Darwin’s abominable mystery, which asks why angiosperms abruptly appeared and rapidly accelerated in their diversification during the Cretaceous period (Friedman, 2009). While more research is required to confirm the lack of subgenome dominance in other plants, fish, and amphibians, the phenomenon has clear affects in agronomic traits (Schnable and Freeling, 2011). Bc genome dominance in *B. carinata* was clearly evident by its increased gene retention, general expression dominance, and lower TE load, with possible effects on biological function (Edger et al., 2017). Additional evidence demonstrated that the Cc subgenome had undergone higher levels of positive selection, whereas the Bc subgenome showed likely purifying selection after polyploidization.

In conclusion, the high-quality *B. carinata* genome reported here will help to improve our comprehension of the genetics underlying favorable agronomic traits. We also observed evidence of the early stages of polyploidization and diploidization-type phenomena that will contribute to our understanding of interspecific hybridization and the establishment of subgenome dominance in *Brassica*. Our identification of Bc subgenome dominance in the *B. carinata* genome will significantly aid efforts in its improvement as a biofuel resource. Efforts for crop improvement, such as addressing the plant’s tall stature and long life cycle, have been thwarted by the species’ lack of genetic diversity (Sheikh et al., 2014). Our comprehensive characterization of the *B. carinata* could advance crop improvement efforts, including those of other *Brassica spp.* through introgression. The addition of this excellent *B. carinata* genome assembly to complete the *Triangle of U* will further accelerate the pace of genetic and genomic research in *Brassica*.

## Methods

### Genome assembly and annotation

The *B. carinata* genome was assembled from PacBio sequencing data using the Hi-C scaffolding method. Contigs were assembled using Canu version 2.0 (Koren et al., 2017). A reference-assisted method was applied to cluster the contigs in reference to their respective subgenomes, *B. nigra* (Bn) and *B. oleracea* (Co), using RaGoo (Alonge et al., 2019). The Hi-C data was used to generate scaffolds for 17 pseudomolecules using ALLHIC (Zhang et al., 2019). The scaffolding results were inspected with the 3D-DNA method (Dudchenko et al., 2017), followed by manual correction. The pseudomolecules were consecutively subjected to error correction using Pilon (Walker et al., 2014), Arrow (PacificBiosciences, 2020), and FreeBayes (Garrison and Marth, 2012) with Illumina paired-end reads.

Repetitive elements within the *B. carinata* genome were masked using a series of programs based on the MAKER Advanced Repeat Library Construction documents (Campbell et al., 2014). We used a MAKER pipeline to annotate protein-coding sequences within the genome based on multiple forms of evidence. Briefly, a total of six rounds of MAKER (Cantarel et al., 2008) was run with SNAP (Korf, 2004) and AUGUSTUS (Stanke et al., 2006) for *ab initio* gene prediction with full-length candidates from *Arabidopsis*, *B. nigra*, *B. oleracea*, *B. napus,* and *B. juncea* comprising the protein database. The completeness of our gene annotation was assessed by analyzing the presence of BUSCOs resulting from each round of the MAKER (Cantarel et al., 2008) process. The detailed methods are described in the Supplementary Text.

For functional description annotation, DCBLAST (Yim and Cushman, 2017) was used to identify the homologous genes in the *Arabidopsis* protein database (Krishnakumar et al., 2015), UniProt-SwissProt, and UniProt-TrEMBL (Boutet et al., 2016). The descriptions for each predicted *B. carinata* protein were inferred from the human-readable protein descriptions using AHRD (Schoof, 2020), based upon protein names from the three protein databases. We identified the Pfam domains, Gene Ontology (GO) terms, and Kyoto Encyclopedia of Genes and Genomes (KEGG) pathway information associated with each protein using InterProScan (Jones et al., 2014). We used Pfam domains to identify transcription factors using plantTF_identifier [176]. Pathways were identified using the Ensemble Enzyme Prediction Pipeline (E2P2 v3.1) (E2P2 v3.1 - Ensemble Enzyme Prediction Pipeline version 3.1 | Plant Metabolic Network). For functional enrichment analysis, GO terms from InterProScan were used [175]. A *B. carinata*-specific GO database was created to predict the enrichment of GO terms and also used GOATOOLS (Klopfenstein et al., 2018) for GO term enrichment analysis.

### Transcriptome sequencing sample collection and analysis

Seven cDNA libraries were generated from the following *B. carinata* tissues for sequencing: (1) 13-day-old whole seedlings grown on Murashige and Skoog (MS) media (Murashige and Skoog, 1962); (2) roots from 7-week-old seedlings grown on MS medium; (3) young leaves from 7-week-old plants grown on MS medium; (4) leaves from mature plants grown in soil; (5) stems from mature plants; (6) flowers from mature plants following anthesis; and (7) immature siliques from mature plants. For heat stress analyzes, 18-day-old *B. carinata* seedlings were subjected to 8 h of heat stress. The heat-stressed plants were well watered before the growth chamber temperature was raised to 40 °C at 8:00 am PST, at 2 h past the beginning of the photoperiod. Tissue samples were collected and immediately snap-frozen in liquid N2, and RNA was extracted from the 15 tissue samples as described in the Supplementary Text. RNA-Seq data were obtained as described in the Supplementary Text.

The RNA-Seq data were preprocessed using TrimGalore (Krueger, 2015) to remove any low-quality reads using a minimum Phred-like quality score (Q-score) of 20 and a minimum length of 50 bp, leaving the validated read pairs for subsequent analysis. The RNA-Seq reads were then aligned to the current version of the *B. carinata* annotated genome using the STAR aligner (Dobin et al., 2013). Read counts were obtained using featureCount (Liao et al., 2014) to identify quantitative differential expression estimates using DESeq2 (Love et al., 2014). A false discovery rate (FDR) cutoff of <0.001 and a +/- 2-fold-change was implemented to determine differentially expressed genes (DEGs). The TPM normalization method was used for downstream gene expression analysis (Robinson et al., 2010).

### Comparison of orthologous genes and gene families

To investigate the expansion or contraction of gene clusters, the gene families generated by OrthoFinder (Emms and Kelly, 2019) and the phylogenetic tree structure of 27 species were subjected to a computational analysis of changes in gene family sizes using a CAFE (Computational Analysis of gene Family Evolution) calculation (version 2.1) (Han et al., 2013). We also performed GO term enrichment and KEGG pathway analyzes were also performed, OmicShare (OmicShare). Gene duplications were identified using the DupGen_finder pipeline (Qiao et al., 2019), with the DupGen_finder-unique.pl script and were classified into different gene duplication types. The diploid progenitors were taken as outgroups for all species in the *Triangle of U*. A genome wide BLASTP search was performed (e-value < 1e−10, maximum 5 matches, and tabular format) using the DCBLAST (Yim and Cushman, 2017) pipeline followed by MCScanX (Wang et al., 2012) analysis to identify homoeolog pairs and then to classify the gene duplications as tandem, proximal, transposed, or dispersed. To identify proteins targeted by pathogens, we explored variable domain architectures were explored in *Triangle of U* species using plant_rgene (Sarris et al., 2016). The domains present in the predicted proteins from each species were identified using the Pfam database (Finn et al., 2014) using HMMER (Mistry et al., 2013). The domain annotations were parsed, and NB-ARC domain-containing proteins were identified and analyzed for the presence of additional fused domains using the plant_rgene pipeline (Sarris et al., 2016).

### Dating of whole genome duplication and genome and species divergence

Molecular dating was carried out using a stringent set of 1,181 low-copy orthologues (411,367 sites) in all the *Brassica* subgenomes (An, Ar, Aj, Bc, Bn, Bj, Cc, Cn, and Co), *Arabidopsis thaliana*, *Arabidopsis lyrata*, *Arabis alpina*, and *Raphanus sativus*, together with six fossil-based age constraints on internal nodes of the tree. Both the peptide sequences and coding sequences (CDSs) of these orthologue groups were aligned using MUSCLE (Edgar, 2004). The CDSs were aligned onto the amino acid alignments using PAL2NAL (Suyama et al., 2006). Any invalid alignments were then filtered out using the following trimAI criteria (Capella-Gutiérrez et al., 2009). If there were gaps in over 90% of the sequences, sequence bases in the alignments were removed. Second, if the translations of the transcript coverage comprised less than 30% of the total alignment length of a gene family, they were also filtered out. A maximum likelihood (ML) analysis was performed with the 1,181-sequence supermatrix using RaxML (Stamatakis, 2014) under the GTR + GAMMA + I model defined as the best-fit evolutionary model according to the Bayesian information criterion (BIC) from IQ-TREE (Nguyen et al., 2015). We calculated the posterior probability distribution of node ages and evolutionary rates were calculated using a Markov chain Monte Carlo (MCMC) tree in the PAML package (Yang, 1997).

We performed local gene collinearity analysis to identify homoeologs in *B. carinata* (BcCc), *B. juncea* (AjBj), *B. napus* (AnCn), *B. nigra* (Bn), and *B. oleracea* (Co). We performed an all-against-all sequence similarity analysis using LAST (LAST: genome-scale sequence comparison). Collinearity blocks were identified for at least four homoeologs pairs with a distance cutoff of 20 genes with an analysis processed by MCScanX (Wang et al., 2012). For each of the resulting homoeologous pairs, we then calculated synonymous substitutions per synonymous site (*Ks*) using yn00 in the PAML package (Yang, 1997). The *K*s distribution peaks for each species were identified using Gaussian Mixture Models (GMM) with the Dupgene pipeline (Qiao et al., 2019).

The hybridization date for a given *B. carinata* genome was estimated from the average of two calculations: *KsB*/2TB and *KsC*/2TC, where *KsB* is the *Ks* rate estimated for ortholog pairs in the *B. nigra* and Bc subgenomes, *KsC* is the *Ks* rate estimated for ortholog pairs in a given *B. oleracea* and Cc subgenome, and *T* is the divergence time proposed in previous reports (Lysak et al., 2005; Navabi et al., 2013; Arias et al., 2014; Cheng et al., 2017). For instance, *B. carinat adivtime = KsB(Bni,BcaB)2 TBBni+ KsC(Bol,BcaC)2 TCBol*. *KsB* and *KsC* were determined as the mean *Ks* value from all orthologous pairs analyzed.

## Accession Numbers

Sequence data from this article can be found in the EMBL/GenBank data libraries under accession number(s): PacBio (SRRXXXXXX), Hi-C (SRRXXXXXX), RNA-Seq (SRRXXXXXX)

## Supplemental Data

Supplemental Figure 1. Morphological comparison of *B. carinata* organs from different growth stages.

Supplemental Figure 2. PacBio sequencing data summary.

Supplemental Figure 3. Paired-end read duplication levels for the Illumina sequencing reads.

Supplemental Figure 4. HiC-Pro quality metrics for Hi-C data.

Supplementary Figure 5. The genome sequencing and reference-based assembly of *Brassica carinata*.

Supplementary Figure 6. Insertion time of intact Long terminal repeat retrotransposons (LTR-RTs) groups in *Brassica* whole genome species.

Supplementary Figure 7. Transcriptome cDNA read statistics.

Supplementary Figure 8. Genome annotation.

Supplementary Figure 9. Quantitative assessment of Brassicaceae genomes and transcriptomes using BUSCO.

Supplementary Figure 10. BUSCO assessment of genome assembly completeness among the *Triangle of U* (sub)genomes.

Supplementary Figure 11. Gene map of the *B. carinata* chloroplast genome.

Supplementary Figure 12. KEGG pathway enrichment analysis of expanded gene families among *Triangle of U* species.

Supplementary Figure 13. Top 20 enriched KEGG pathways among the six *Triangle of U Brassica* species.

Supplementary Figure 14. Top 20 enriched Gene Ontology (GO) terms in the biological process domain among the *Triangle of U* species.

Supplementary Figure 15. Top 20 enriched Gene Ontology (GO) terms in the cellular component domain among the *Triangle of U* species.

Supplementary Figure 16. Top 20 enriched Gene Ontology (GO) terms in the molecular function domain among the *Triangle of U* species.

Supplementary Figure 17. Transcription factor families identified in the *B. carinata* genome.

Supplementary Figure 18. Assessment of biosynthetic and metabolite pathway completeness in the *B. carinata* genome.

Supplementary Figure 19. Phylogenetic tree of FLC homologs from *Brassica* species.

Supplementary Figure 20. Expression profiles of 13 fatty acids from the seeds of seven *B. carinata* accessions.

Supplementary Figure 21. PCA plots of seed fatty acid contents of seven *B. carinata* accessions.

Supplementary Figure 22. Phylogenetic tree of *Triangle of U* species homologs encoding lipid transfer protein.

Supplementary Figure 23. Phylogenetic tree of phenylalanine ammonia-lyase 1 homologs from *Triangle of U* species.

Supplementary Figure 24. Expression profiles of eight glucosinolate metabolites from the seeds of seven *B. carinata* accessions.

Supplementary Figure 25. Phylogenetic tree of TGG4a homologs from *Brassica* species.

Supplementary Figure 26. Syntenic dot plot intragenomic comparison of the *B. carinata* genome assembly.

Supplementary Figure 27. Top 20 Gene Ontology (GO) terms for proximal duplicates among the *Triangle of U* genomes.

Supplementary Figure 28. Top 20 Gene Ontology (GO) terms for tandem duplicates among the *Triangle of U* genomes.

Supplementary Figure 29. Distribution of genomic features among the subgenomes of the *Triangle of U* allotetraploid species.

Supplementary Figure 30. Typical homoeologous exchanges in *B. carinata*.

Supplementary Figure 31. Pattern of gene expression dominance and relative selection pressure between homoeologous gene pairs.

Supplementary Figure 32. Identification of collinearity between the *B. carinata* subgenomes.

Supplementary Figure 33. Bc subgenome homoeologous exchanges in *B. carinata*.

Supplementary Figure 34. Cc subgenome homoeologous exchanges in *B. carinata*.

Supplementary Figure 35. Homoeologous exchanges between *B. carinata* chromosomes and those of its diploid progenitor genomes.

Supplementary Figure 36. Syntenic dot plots of alignment of the *Brassica carinata* var. Gomenzer genome against *Brassica carinata* var. Zd-1 genome.

Supplementary Table 1. Flow cytometry genome size estimation for *B. carinata*.

Supplementary Table 2. PacBio sequencing read summary from the SMRT portal.

Supplementary Table 3. Illumina sequencing read summary from MultiQC.

Supplementary Table 4. Draft genome scaffolding results during the assembly and error correction phases.

Supplementary Table 5. Error correction results.

Supplementary Table 6. Scaffolding results of the draft genome assembly.

Supplementary Table 7. Flow cytometry genome size estimations among *Triangle of U* assemblies.

Supplementary Table 8. Summary of repeat analysis of the B. carinata genome assembly.

Supplementary Table 9. Repeat content per chromososome of the *B. carinata* genome.

Supplementary Table 10. Summary of repeat analysis among the *Triangle of U* genomes.

Supplementary Table 11. Evaluation of LTR Assembly Index (LAI) values among the *Triangle of U* (sub)genomes.

Supplementary Table 12. Contig results of the de novo transcriptome assembly for B. carinata.

Supplementary Table 13. Gene annotation statistics among B. carinata and related *Triangle of U* (sub)genomes.

Supplementary Table 14. Mapping and alignment statistics of the RNA-Seq data.

Supplementary Table 15. Transposable element related gene models.

Supplementary Table 16. Coding sequence annotation statistics for the B. carinata genome.

Supplementary Table 17. The synonymous base substitution (Ks) and divergence time estimations among *Triangle of U* species and radish.

Supplementary Table 18. Expansions and contractions of gene families among the *Triangle of U* species.

Supplementary Table 19. Size distribution of transcription factor families among B, carinata and shared *Triangle of U* (sub)genomes.

Supplementary Table 20. *Arabidopsis* genes related to flowering time and their orthologs in the *Triangle of U* genomes.

Supplementary Table 21. *Arabidopsis* genes related to acyl lipid metabolism and their orthologs in the *Triangle of U* genomes.

Supplementary Table 22. *Arabidopsis* genes related to anthocyanin metabolism and their orthologs in the *Triangle of U* genomes.

Supplementary Table 23. *Arabidopsis* genes related to glucosinolate metabolism and their orthologs in the *Triangle of U* genomes.

Supplementary Table 24. Survey of nucleotide-binding site (NBS) proteins among the *Triangle of U* (sub)genomes.

Supplementary Table 25. Summary of orthologs in the *Triangle of U* species and *A. thaliana*.

Supplementary Table 26. List of collinear blocks between *B. carinata* and its progenitor species, *B. nigra* and *B. oleracea*.

Supplementary Table 27. Duplicate gene analysis in *B. carinata* and shared *Triangle of U* (sub)genomes.

Supplementary Table 28. Sequence evolution rates of duplicate genes in *B. carinata* and the other *Triangle of U* genome assemblies.

Supplementary Table 29. Survey of paralogs in the *B. carinata* genome.

Supplementary Table 30. Confirmed genes deleted in *B. carinata* and shared *Triangle of U* (sub)genomes.

Supplementary Table 31. Homoeologous gene sets among *B. carinata* subgenomes and their respective progenitor genomes (Bn and Co).

Supplementary Table 32. Summary of *Ka* and *Ks* values among *B. carinata* and its progenitor species, *B. nigra* and *B. oleracea*.

Supplementary Table 33. Position of homoeologous exchange events and bootstrap values for of Maximum Likelihood (ML) tree construction.

Supplementary Table 34. Position of homoeologous exchange events (Bc gene replaced) and bootstrap values for of Maximum Likelihood (ML) tree construction.

Supplementary Table 35. Position of homoeologous exchange events (Cc homoeolog replaced) and bootstrap values for of Maximum Likelihood (ML) tree construction.

Supplementary Table 36. Position of homoeologous exchange events (Bc homoeolog lost) and bootstrap values for of Maximum Likelihood (ML) tree construction.

Supplementary Table 37. Position of homoeologous exchange events (Cc homoeolog lost) and bootstrap values for of Maximum Likelihood (ML) tree construction.

Supplementary Table 38. Gene Ontology enrichment analysis of homoeologous exchanges in *B. carinata*.

Supplementary Table 39. Homoeolog and paralog gene expression comparisons between *B. carinata* subgenomes.

Supplementary Table 40. Transposon insertion locations and transcript abundance of *FAE1* in the *B. carinata* genome.

Supplementary Table 41. Transposon insertion locations and transcript abundance of *FLC* in the *B. carinata* genome.

Supplementary Table 42. Transposon insertion locations and transcript abundance of *FAR1* in the *B. carinata* genome.

Supplementary Table 43. Isolated glucosinolates in seven *B. carinata* accessions.

Supplementary Table 44. Subcellular localization of *B. carinata* genes.

Supplementary Table 45. Automated Assignment of Human Readable Descriptions (AAHRD) results for *B. carinata* genes.

Supplementary table 48. *Arabidopsis* genes related to mediating the auxin response and their orthologs in the *Triangle of U* genomes.

## Acknowledgements

We gratefully acknowledge the support of the Nevada Agricultural Experiment Station (Grant No. NEV00384) and VPRI research funding (University of Nevada, Reno). The Pires lab is funded by the National Science Foundation (NSF IOS 1339156) and the Department of Energy Defense Threat Reduction Agency (HDTRA 1-16-1-0048). The Edger lab is funded by the National Science Foundation (NSF IOS 2029959). The Mason lab is partially funded by the Deutsche Forschungsgemeinschaft (DFG, German Research Foundation) under Germany’s Excellence Strategy (EXC 2070 - 390732324). The authors would like to thank the Germplasm Resources Information Network of the USDA-ARS for their help in providing genetic resources. The authors would also like to thank Maggie Weitzman at the Genomics & Cell Characterization Core Facility (GC3F) at the University of Oregon for performing the PacBio library preparation and sequencing. Further thanks go to Diana Burkart-Waco for carrying out the Illumina sequencing at the DNA Technologies and Expression Analysis Cores at the UC Davis Genome Center, supported by NIH Shared Instrumentation (NIH 1S10OD010786-01). We would also like to thank Sitharam Ramaswami at the Genome Technology Center at NYU Langone Health for performing Hi-C sequencing. Finally, we would like to thank the Office of Information Technology and Research & Innovation at the University of Nevada, Reno providing paid access to the Pronghorn High-Performance Computing Cluster.

## Author Contributions

WCY led and managed the project. HDH, MLS, and WCY collected, prepared, and sequenced the plant material. ASM, PE, JCC, JH, JCP, JS, HT, WCY, and XZ devised the main conceptual ideas and proof outline. WCY, HT, and XZ assembled and annotated the genome. DM, SW, WCY, and XZ performed gene family and gene enrichment analysis. DDC and WCY performed Hi-C analysis. ALN, DAP, and WCY performed comparative evolutionary analysis. ASM, HA, KB, PE, JCP, HT, and WCY designed and performed homoeologous exchange analysis. All authors contributed to the article and approved the submitted version.

## Corresponding Author

Correspondence to Won Cheol Yim

## Author information Affiliation

University of Nevada, Reno, NV, USA

Won Cheol Yim, Mia L. Swain, Hyun Don Ham, David D. Curdie, Agusto Luzuriaga-Neira, Samuel Wang, Juan K. Q. Solomon, Jeffrey F. Harper (jharper.unr.edu), Dylan K. Kosma, David Alvarez-Ponce, John C. Cushman

Fujian Agriculture and Forestry University, Fuzhou, China Dongna Ma, Haibao Tang, Xingtan Zhang University of Missouri, Columbia, Columbia, MO, United States Hong An, J. Chris Pires Michigan State University, East Lansing, MI, United States

Kevin A. Bird, Patrick P. Edger

University of California, Riverside, Riverside, CA, United States Jay S. Kirkwood (jkirk.ucr.edu), Manhoi Hur The University of Bonn, Bonn, Germany

Annaliese S. Mason

